# MED1 IDR acetylation reorganizes the transcription preinitiation complex, rewires 3D chromatin interactions and reprograms gene expression

**DOI:** 10.1101/2024.03.18.585606

**Authors:** Ran Lin, Douglas Barrows, Yan Mo, Takashi Onikubo, Zhiguo Zhang, Robert G. Roeder

## Abstract

With our current appreciation of the complexity of eukaryotic transcription, whose dysregulation drives diseases including cancer, it is becoming apparent that identification of key events coordinating multiple aspects of transcriptional regulation is of special importance. To elucidate how assembly of RNA polymerase II (Pol II) with Mediator complex preinitiation complexes (PICs) and formation of transcription-permissive 3D chromatin organization are coordinated, we studied MED1, a representative subunit of the Mediator complex that acts to establish functional preinitiation complexes (PICs)^1^ that forms biomolecular condensates through an intrinsically disordered region (IDR) to facilitate transcription^2^, and is implicated in the function of estrogen receptor α (hereafter ER) in ER-positive breast cancer (ER^+^ BC) cells^3,4^. We found that MED1 is acetylated at 6 lysines in its IDR and, further, that MCF7 ER^+^ BC cells in which endogenous MED1 is replaced by an ectopic 6KR (non-acetylatable) mutant (6KR cells) exhibit enhanced cell growth and elevated expression of MED1-dependent genes. These results indicate an enhanced function of 6KR MED1 that may be attributed to two mechanisms: (1) reorganized PIC assembly, as indicated by increased MED1 and Pol II, decreased MED17, and equivalent ERα occupancies on chromatin, particularly at active enhancers and promoters; (2) sub-TAD chromatin unfolding, as revealed by HiCAR (Hi-C on accessible regulatory DNA) analyses. Furthermore, in vitro assays demonstrate distinct physio-chemical properties of liquid-liquid phase separation (LLPS) for 6KR versus 6KQ MED1 IDRs, and for non-acetylated versus CBP-acetylated WT MED1 IDR fragments. Related, Pol II CTD heptads are sequestered in 6KR and control WT MED1 IDR condensates, but not 6KQ and CBP-acetylated WT MED1 IDR condensates. These findings, in conjunction with recent reports of PIC structures^5–7^, indicate that MED1 coordinates reorganization of the PIC machinery and the rewiring of regional chromatin organization through acetylation of its IDR. This study leads to an understanding of how the transition in phase behavior of a transcription cofactor acts as a mechanistic hub integrating linear and spatial chromatin functions to support gene expression, and have potential therapeutic implications for diseases involving MED1/Mediator-mediated transcription control.

## Introduction

In eukaryotic cells, all protein-coding genes and many non-coding RNAs are transcribed by RNA Pol II in conjunction with the Mediator complex. Upon recruitment by transcription factors (TFs), the 26-subunit 1.4 MDa Mediator complex acts to establish functional preinitiation complexes (PICs) at core promoters of target genes to initiate transcription^1,8^. In support of the essential function of Mediator in transcription, cryo-EM structural studies have documented dynamic interactions between Mediator and Pol II in human cells^5–7^. Acting at this dynamic interface, the Mediator MED1 subunit forms a flexible Rod-Elbow tether with MED4 and MED9 subunits^5,6^.

The 3D chromatin structure is heralded as an important factor in the regulation of gene expression, but the function of Mediator and Pol II in 3D chromatin organization remains a matter of active debate. In contrast to the limited effects observed by traditional chromatin capture assays^9–11^, recent studies employing single nucleosome resolution micro-C assays^12^ have revealed critical roles of Mediator^13^ and Pol II^14^ in enhancer-promoter interactions and formation of CTCF-dependent loops. Moreover, Mediator (through its kinase module) restricts the spread of chromatin compaction and is therefore required for maintenance of chromatin domains permissible for gene expression^15^, corroborating an earlier report that gene activation in yeast occurs on chromatin with less compaction^16^.

IDRs are low-complexity fragments of proteins that support dynamic multi-valent interactions and LLPS of proteins^17^, and are over-represented in transcription factors and cofactors^8^. Although the impact of the large MED1 IDR segment on the cryo-EM structure of Mediator has not been precisely determined^5^, this IDR promotes condensation of MED1 at super-enhancers of genes and corresponding MED1 condensates sequester Pol II^2^ as well as elongation factors CTR9 and SPT6^18^ to facilitate transcription. In addition, various transcription factors^19–21^ and cofactors^22^ have been reported to form nuclear condensates that enable regulation of 3D chromatin organization and gene expression through mechanical forces called chromatin filter^23^, raising the question as to whether Mediator, through its IDR, coordinates PIC assembly and 3D chromatin architecture during gene activation.

More than two-thirds of breast cancers are positive for estrogen receptor α^24^ and over 3000 genes involved in a plethora of tumor-related processes are regulated by ER in BC cells^25^. Interaction of ER with the MED1-containing Mediator complex promotes its function in activating target genes^3,4,26^. MED1 is also over-expressed in a high proportion (40–50%) of primary breast cancers ^27^, further indicative of its important role in ER^+^ BC.

Here we show that the MED1 IDR can be acetylated at six lysines and that the function of a non-acetylatable (6KR) mutant MED1 in ER+ BC cells results in enhanced cell growth and elevated expression of MED1-dependent genes. Genome wide analyses of MED1, MED17 and Pol II occupancies further indicate a 6KR MED1-mediated PIC reorganization and HiCAR analyses indicate that 6KR MED1 rewires 3D chromatin organization by reducing sub-TAD 3D chromatin interactions. The demonstration of distinct physio-chemical properties of LLPS of 6KR versus 6KQ MED1 IDR fragments, and of non-acetylated versus CBP-acetylated WT MED1 IDR fragments, suggests that the 6KR MED1-mediated Pol II recruitment and regional chromatin reorganizations might both be caused by a transition in MED1 phase behavior. Such a concerted regulation of linear and spatial chromatin functions in gene activation suggests that Mediator could integrate multiple mechanisms to achieve gene expression. Targeting such a coordination hub is an inviting therapeutic option for diseases, such as ER^+^ breast cancer, involving MED1/Mediator-mediated transcription.

## Results

### MED1 has an essential function in (MCF7) ER^+^ BC cells

To corroborate the essential role of MED1 in ER target gene transcription, we examined the effects of MED1 depletion in ER^+^ BC cell lines, MCF7 and T47D. MED1 was efficiently depleted by a CRISPR/Cas9 approach with gRNAs targeting 3 independent regions (designated KO-1, −2, and −3) in exons of the *MED1* gene, with a gRNA targeting the mouse *Rosa26* locus as a non-targeting control (Supplementary Fig. 1a, b). An analysis of ER abundance revealed no significant alteration in expression (Supplementary Fig. 1b). As revealed by growth curves, MED1 depletion attenuates cell growth in both MCF7 and T47D cells (Supplementary Fig. 1c). Colony formation for *MED1* KO MCF7 cells is also attenuated (Supplementary Fig. 1d).

Next, sequencing of Poly(A) RNA (RNA-seq) was employed to measure mature RNA expression in *MED1* KO MCF7 cells. Differentially expressed genes (DEGs) from a linear model to characterize the combined effect of KO-1, −2, and −3 were identified and are shown in a volcano plot (Fig. 1a). The differences in the gene expression profiles are also presented as heatmaps (Fig. 1b). An over-representation analysis using a Fisher’s exact test with KEGG (Kyoto Encyclopedia of Genes and Genomes) gene lists revealed that genes involved in cell cycle and DNA replication are enriched in the down-regulated DEGs, underscoring the importance of MED1 for optimal cell growth. In support of a significant role for MED1 in orchestrating ER-driven gene expression, the estrogen signaling pathway is also enriched in down-regulated DEGs. In contrast, analysis for the up-regulated DEGs found various signaling pathways with roles in cell growth, including ErbB (EGFR) and mTOR pathways (Fig. 1c).

**Fig. 1.**
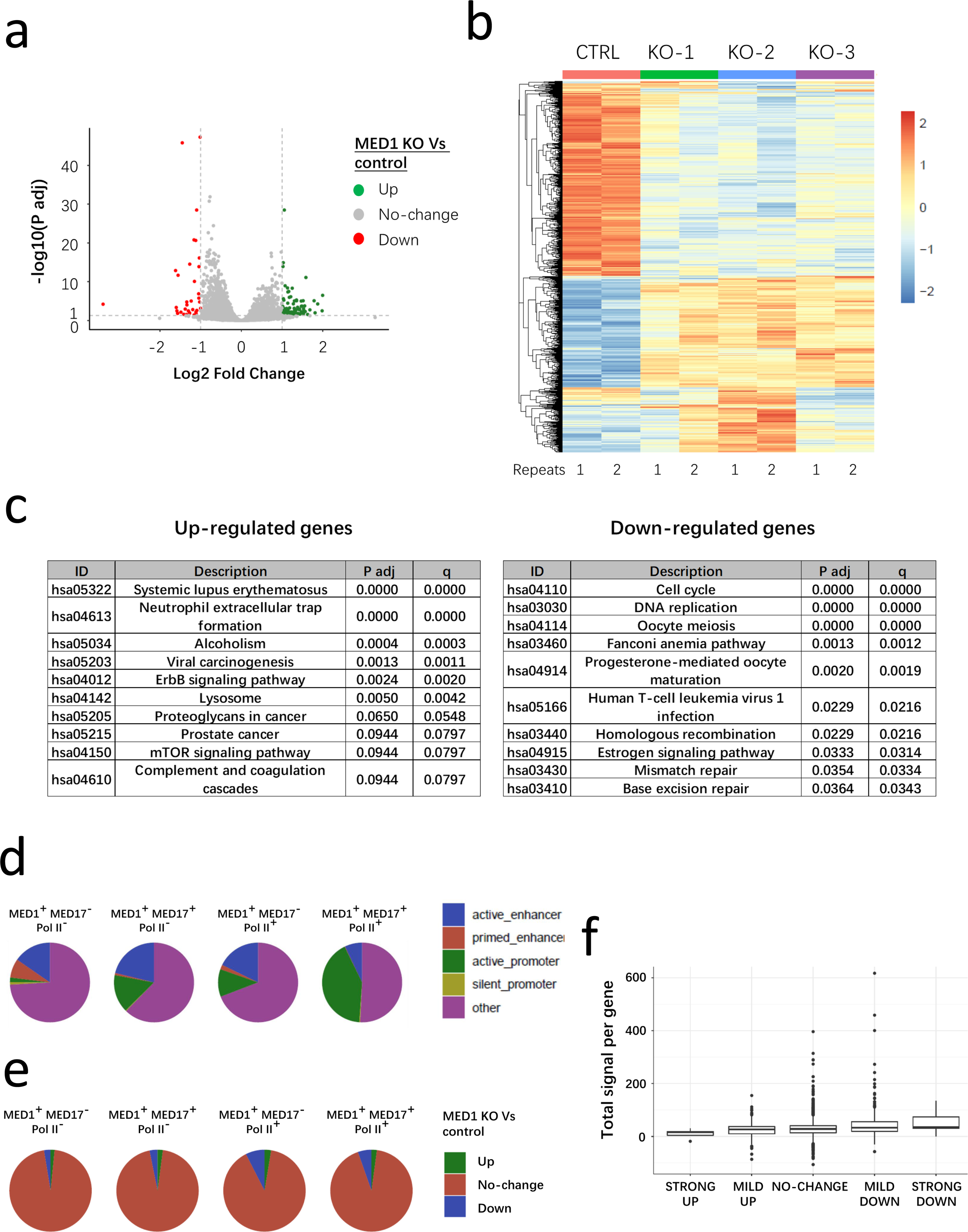
MED1 supports gene activation in MCF7 cells. (a-c) Poly(A) RNA-seq for control and *MED1* KO MCF7 cells. (a) A volcano plot showing the adjusted P value and fold-change of each detected gene (combined from KO−1, −2, and −3) where the cut-off to define DEGs (FC = 2, P adj = 0.05) is indicated. (b) A clustered heatmap of Z-scores of genes with differential expression with P adj < 0.05 from *MED1* KO (−1, −2 and −3) versus control cells. (c) Over-representation analyses using a Fisher’s exact test with KEGG gene lists for up-regulated and down-regulated DEGs, respectively. (d, e) Pie diagrams showing constitution of each type of ChIP-seq peak classified by occupancy of MED1, MED17 and Pol II for types of regulatory elements (d), association with up-, down-regulated genes or no-change-genes of *MED1* KO versus control cells (e). (f) Density of MED1 occupancy at active promoters associated with plotted genes with stratification of genes based on their fold-change values detected from RNA-seq of *MED1* KO Vs control cells.

To further explore the function of MED1 in gene regulation, we elucidated the genomic occupancy of MED1, MED17 and RNA Pol II in MCF7 cells by Chromatin Immunoprecipitation with sequencing (ChIP-seq). As MED17 acts as a scaffolding protein of the core Mediator complex^5,6^, it is reasonable to consider that the presence of MED17 reflects occupancy of the entire core Mediator. Results were corrected for MED1 occupancy arising from non-specific antibody recognition by removing such signals detected in *MED1* KO cells. All MED1-occupied peaks were classified as (a) MED1^+^MED17^−^Pol II^−^ (occupied only by MED1, no apparent occupancy by MED17 or Pol II), (b) MED1^+^MED17^+^Pol II^−^ (co-occupied by MED1 and MED17, no Pol II occupancy), (c) MED1^+^MED17^−^Pol II^+^ (co-occupied by MED1 and Pol II, no apparent MED17 occupancy) and (d) MED1^+^MED17^+^Pol II^+^ (co-occupied by MED1, MED17 and Pol II) peaks. Based on ChIP-seq analysis of histone marks, cis-regulatory elements of genes were assigned as follows: active enhancers: H3K4me^+^ H3K4me3^−^ H3K27ac^+^; primed enhancers: H3K4me^+^ H3K4me3^−^ H3K27ac^−^; active promoters: H3K4me3^+^ within TSS±1kb; silent promoters: H3K4me3^−^ within TSS±1kb^28,29^. Accordingly, MED1-, MED17-, and Pol II-co-occupied peaks were located more at active promoters than enhancers. The MED1^+^MED17^−^ peaks that were detected might reflect the dynamic nature of Mediator-Pol II assembly during transcription activation, as they occupied a larger proportion of primed enhancers than did other peaks (Supplementary Fig. 1d). Next, we examined the association of MED1/MED17/Pol II peaks with DEGs of *MED1* KO cells. To this end, we employed GREAT strategy, in which every gene is assigned a regulatory domain and each genomic region is associated with all genes whose regulatory domains overlap^30^. In contrast to other peaks, MED1- and Pol II-co-occupied peaks (regardless of MED17 occupancy) were associated with more down-regulated DEGs in *MED1* KO cells (Supplementary Fig. 1e). The genes detected by RNA-seq were stratified into 5 tiers (designated STRONG UP, MILD UP, NO-CHANGE, MILD DOWN, STRONG DOWN) according to differential expression in *MED1* KO cells, and MED1 occupancy at active promoters of genes showed the highest value in the tier of STRONG DOWN (Supplementary Fig. 1f). These results lend support to the notion that MED1-containing Mediator is more involved in activation, rather than suppression, of target genes.

### MED1 is acetylated at 6 lysines in the IDR

A large number of transcriptional activators and coactivators have been identified as substrates of acetyl transferases in the past few decades^31^, which prompts the question of whether Mediator subunits are also acetylated in cells. By surveying the acetylation of select Mediator subunits with FLAG tags following ectopic expression in HEK293T cells, we detected acetylation of immunoprecipitated FLAG-MED1 and showed that it was increased both by Trichostatin A (TSA, HDAC inhibitor) and nicotinamide (NAM, Sirtuin inhibitor) (Supplementary Fig. 2a, b). The acetylation of FLAG-MED1 in HEK293T cells was also induced by overexpression of acetyl transferases p300 and CBP, but not by overexpression of a catalytically inactive mutant p300^32^ (Fig. 2a). The p300-induced acetylation of FLAG-MED1 was attenuated by A485, a HAT inhibitor of p300, but not by C646, another HAT inhibitor, or by the bromodomain inhibitor SGC-CBP30 (Fig. 2b). The immunoprecipitated endogenous MED1 also exhibited elevation of acetylation in HEK293T cells overexpressing p300/CBP (Fig. 2c), where the increased levels of MED1 acetylation were correlated with the levels of ectopic p300 (Supplementary Fig. 2c). In contrast to the results with p300 and CBP, ectopic expression of other acetyl transferases that included GCN5, PCAF, TIP60 and MOF did not induce acetylation of FLAG-MED1 in HEK293T cells (Supplementary Fig. 2d). The acetylation of MED1 could be reversed by SIRT1 deacetylase, as ectopically expressed FLAG-MED1 in HEK293T cells exhibited elevated acetylation by treatment of cells with Ex527, a specific SIRT1 inhibitor, in HEK293T cells (Supplementary Fig. 2e). The induction of acetylation of endogenous MED1 by Ex527 was similarly observed in MCF7 cells, but prevented by co-treatment with A485 (Fig. 2d). In addition to Ex527, acetylation of FLAG-MED1 was also induced by Entinostat (inhibitor of HDAC1/2/3) and SAHA (inhibitor of all HDACs) in HEK293T cells (Supplementary Fig. 2f), indicating the potential of both Sirtuin and HDAC type deacetylases in control of MED1 acetylation.

**Fig. 2.**
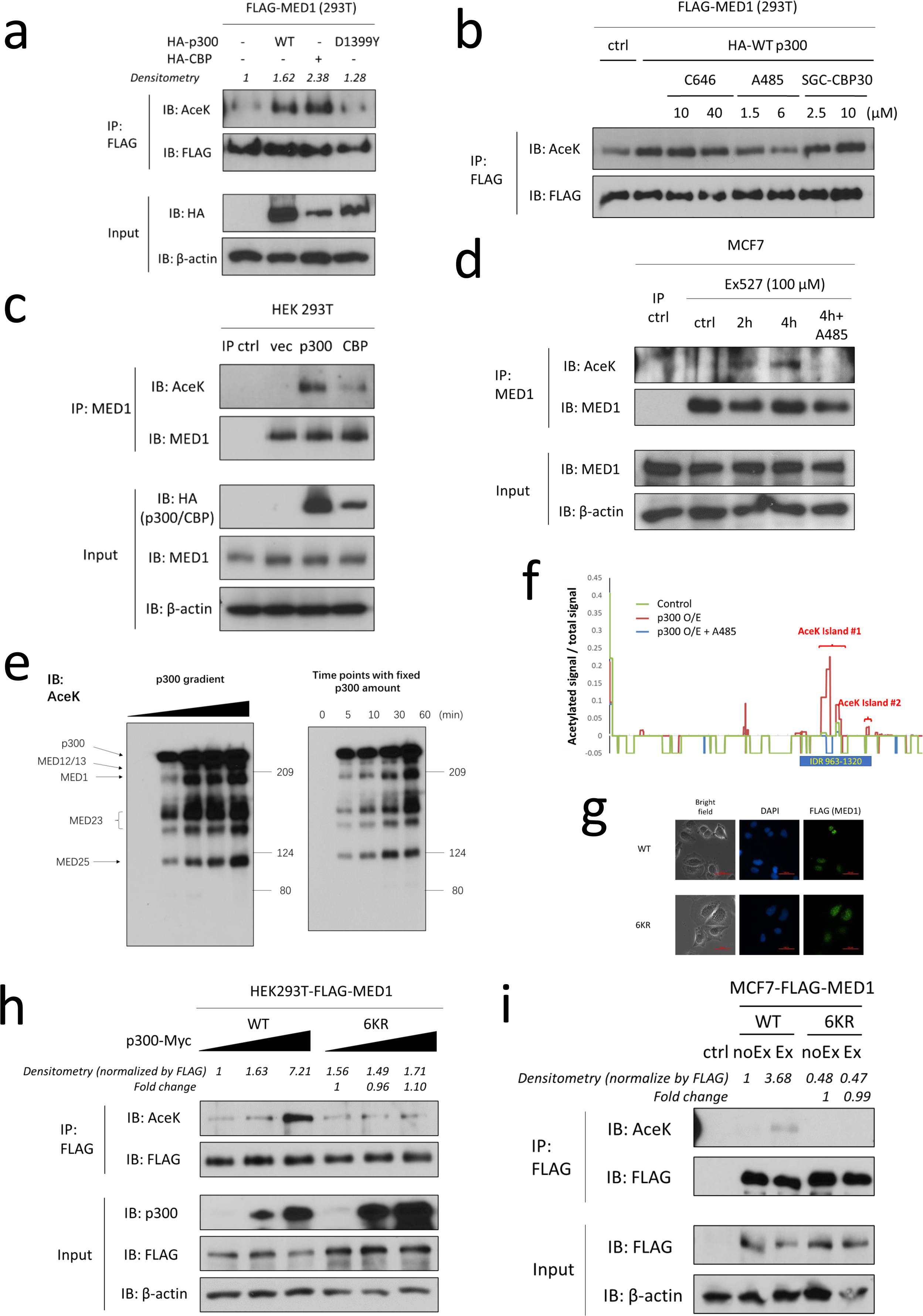
MED1 is acetylated under the control of p300/CBP and SIRT1. (a) Enhanced acetylation (referred as AceK) of ectopically expressed FLAG-MED1 in HEK293T cells following overexpression of p300 (human) or CBP (mouse), but not the catalytically dead p300 D1399Y mutant. (b) Enhanced acetylation of ectopically expressed FLAG-MED1 in HEK293T cells following overexpression of p300 (human) and its suppression by the p300 HAT inhibitor A485, but not by the HAT inhibitor C646 or the bromodomain inhibitor SGC-CBP30. Cells were treated with these inhibitors at the concentrations indicated in the figure for 24 hours. (c) Enhanced acetylation of endogenous MED1 in HEK293T cells following overexpression of p300 (human) or CBP (mouse). IgG was used as an IP control. (d) Enhanced acetylation of endogenous MED1 in MCF7 cells treated with Ex527, an inhibitor of SIRT1, where the effect was attenuated by co-treatment with A485 (6μM, 24 hours). IgG was used as an IP control. (e) Enhanced acetylation of multiple subunits of Mediator (purified from HeLa-S cells) by p300 and acetyl-CoA. Effects of variable p300 concentrations with fixed incubation time (left) and variable incubation times with a fixed amount of p300 (right) were tested. The previously reported auto-acetylation of p300 was also observed. (f) Signal of digested peptides with acetylation plotted as a function of amino acid sequence. Signals from control samples and from p300 overexpression samples with or without A485 were compared and the 2 acetylation clusters are indicated. (g) Immunostaining with anti-FLAG antibodies combined with DAPI staining for MCF7 cells with ectopic expression of FLAG-tagged WT or 6KR MED1. Scale bar is 200 μm. (h) Enhanced acetylation of ectopically expressed WT, but not 6KR, FLAG-MED1 in HEK293T cells following overexpression of p300. Effects of variable p300 concentrations were tested. (i) Enhanced acetylation of ectopically expressed WT, but not 6KR, FLAG-MED1 in MCF7 cells treated with Ex527 (100μM, 2/4 hours, referred as Ex). A control sample was included in which lysate of normal MCF7 cells was immunoprecipitated with anti-FLAG. For (a-d) and (h-i), acetylation of immunoprecipitated FLAG-MED1 or endogenous MED1 was monitored by immunoblotting with pan-anti-acetyl-lysine antibody as described in Methods. The AceK immunoblot signals were normalized by the FLAG signals where densitometry is shown.

The acetylation of MED1 by p300/CBP was also examined in an in vitro assay. The Mediator was purified with anti-FLAG antibodies from nuclear extract of HeLa-S cells expressing FLAG-tagged MED10, a core Mediator subunit. Immunoblot analyses confirmed the integrity of purified Mediator, by monitoring MED1 and MED17, as well as the absence of p300 and SIRT1 (Supplementary Fig. 2g). Incubation of the Mediator complex with full length p300 and acetyl-CoA resulted in acetylation of multiple Mediator subunits, including MED1, MED23 and MED25 in the tail module, and MED12 and MED13 in the kinase module. The previously described auto-acetylation of p300 was also evident. The levels of acetylation of the Mediator subunits and p300 were dependent on the duration and amounts of p300 treatment (Fig. 2e).

Indicative of a dependence of MED1 acetylation on its interaction with nuclear receptors such as ERα, a FLAG-tagged MED1 bearing deletions of the NR boxes that determine strong MED1-nuclear receptor interactions^33^ exhibited attenuated induction of acetylation by Ex527 in MCF7 cells (Supplementary Fig. 2h). The functional importance of estrogen signaling in ER+ BC cells prompted us to investigate its role in MED1 acetylation. The ER agonist 4-Hydroxytamoxifen (4-OHT), which blocks estrogen binding, and Fulvestrant, an ER antagonist that blocks ER-mediated transcription and induces ER degradation, both significantly reduced acetylation of FLAG-tagged MED1 in MCF7 cells (Supplementary Fig. 2i). Therefore, estrogen signaling either directly affects MED1 acetylation or helps to establish conditions favorable for MED1 acetylation.

In order to identify the acetylated lysine residues in MED1, we employed a Liquid Chromatography-Mass Spectrometry (LC-MS)/MS workflow to examine immunoprecipitated endogenous MED1 from HEK293T cells with no treatment (control) or with p300 overexpression with or without A485 (Supplementary Fig. 3a, b). Acetylation levels at K1076, K1113, and K1152 (referred as “island #1”) and at nearby K1300, K1309, and K1311 (referred as “island #2”) were appreciably higher in p300-overexpressing cells than in no-treatment or A485-treated control cells (Fig. 2f). It is noteworthy that these six lysines are located in a 963-1320 residue MED1 fragment that is predicted to be an IDR by the VLXT algorithm^34^ (Supplementary Fig. 3c), as was noted previously^2^. Regarding the importance of IDR-mediated LLPS of MED1 in transcriptional regulation^2,18^ these six lysines became the focus of our study. The vast majority of FLAG-tagged MED1, either WT or the non-acetylatable 6KR mutant (with six lysines replaced by arginines), is localized in cell nuclei in MCF7 cells (Fig. 2g), excluding a scenario in which these six lysines control MED1 function via nuclear entry. In HEK293T cells, p300 alone induced robust acetylation of WT FLAG-MED1, but showed only a limited effect on 6KR FLAG-MED1 acetylation (Fig. 2h). Similarly, Ex527 alone induced acetylation of WT FLAG-MED1 in MCF7 cells, but showed no effect on 6KR FLAG-MED1 acetylation (Fig. 2i). The indicated six lysines are positioned as conserved residues by sequence alignments of MED1 from various species, indicative of their evolutionary significance in MED1 (Supplementary Fig. 3d). In summary, our results identify six major acetylation sites in the MED1 IDR and further show that these acetylation events are regulated p300/CBP and SIRT1.

### The MED1 acetylation mutant affects cell growth and gene expression in ER^+^ BC cells

As the 6 conserved lysines in the MED1 IDR were identified as acetylation substrates, we next sought to determine the functional significance of MED1 acetylation in ER^+^ BC cells. Using CRISPR/Cas9, endogenous MED1 was depleted in a pool of MCF7 cells that ectopically expressed CRISPR-resistant WT or non-acetylatable 6KR MED1 (Supplementary Fig. 4a). In these cells, the ectopic HA-tagged WT and 6KR MED1 proteins replaced the endogenous MED1 with similar levels of expression, and ERα expression showed no significant alteration (Supplementary Fig. 4b). To avoid selection and expansion of advantageous clones from the pool, which would change the expression level of MED1, all experiments were conducted with cells collected immediately after the knock-out of *MED1* by lentiviral infection. By monitoring cell viability, we found that 6KR MED1 add-back MCF7 cells (referred as 6KR/KO cells) exhibit an elevated growth rate compared with WT MED1 add-back MCF7 cells (referred as WT/KO cells) (Fig. 3a, Supplementary Fig. 4c). In parallel, colony formation assays showed a higher growth capability of 6KR/KO cells relative to WT/KO cells (Fig. 3b). Therefore, the non-acetylatable 6KR MED1 enhances the growth of MCF7 cells in culture, indicative of an anti-tumor role of MED1 acetylation in ER^+^ breast cancer.

**Fig. 3.**
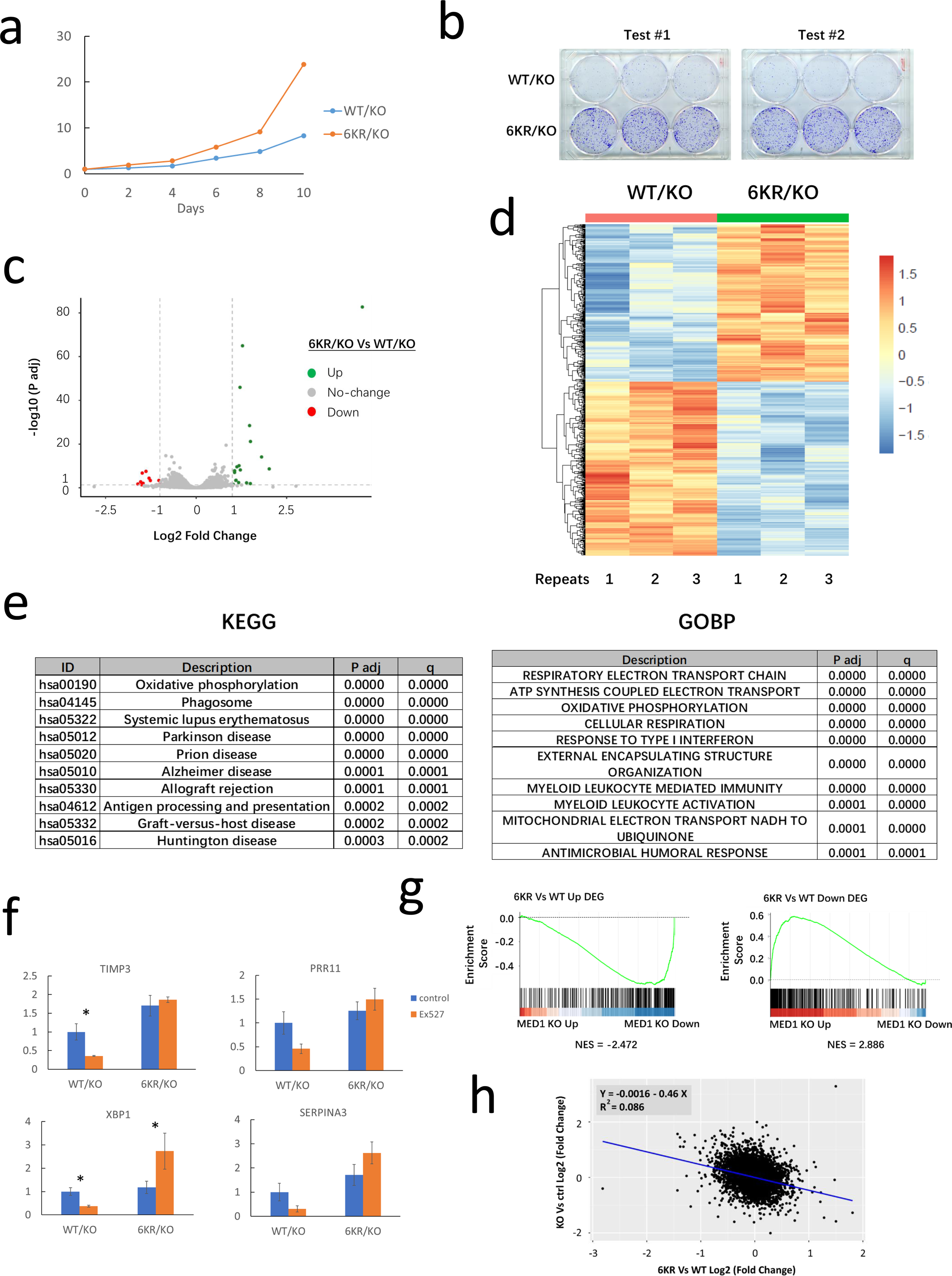
The non-acetylatable 6KR mutation of MED1 enhances cell growth and MED1 target gene expression in MCF7 cells. (a) Elevated growth rate of 6KR/KO versus WT/KO MCF7 cells monitored by CellTiter Blue assay. (b) More colonies formed by 6KR/KO versus WT/KO MCF7 cells stained with crystal violet following growth for 10 days after initial seeding. Two repeat tests are presented. (c-e) Poly(A) RNA-seq for 6KR/KO and WT/KO MCF7 cells. (c) Volcano plot presentation of the adjusted P value and fold-change for each detected gene, with the cut-off for defining DEGs (FC = 2, P adj = 0.05) indicated. (d) A clustered heatmap of Z-scores of genes with differential expression with P adj < 0.05 from 6KR/KO versus WT/KO cells. (e) GSEA analyses comparing the 6KR/KO versus WT/KO gene rank list with KEGG and GOBP gene lists. (f) Expression of select DEGs monitored by qRT-PCR in WT/KO and 6KR/KO cells with and without Ex527 treatment (100 μM, 3.33 hours). N = 3 for each group. Bars report mean ± SE. Statistical significance for control versus Ex527 groups was shown. * p < 0.5 (t test). (g) GSEA analysis showing that the up-regulated (left) and down-regulated (right) genes in 6KR/KO versus WT/KO cells were enriched in the set of down-regulated and up-regulated genes of *MED1* KO (versus control) cells, respectively. (h) Fold-change values of each detected gene from *MED1* KO versus control plotted as a function of fold-change values from 6KR/KO versus WT/KO. The data show an inverse correlation. Each dot represents one gene.

To investigate the ability of MED1 acetylation to regulate gene expression, we performed poly(A) RNA-seq to measure mature RNA expression in 6KR/KO and WT/KO MCF7 cells. The results are shown with a volcano plot (Fig. 3c) and DEGs are shown with a heatmap (Fig. 3d). GSEA (Gene Set Enrichment Analysis) analyses comparing the 6KR/KO versus WT/KO gene rank list with KEGG and Gene Ontology Biological Processes (GOBP) gene lists showed that mitochondrial respiration-related genes are enriched, indicating a mechanism by which 6KR MED1 alters grown of MCF7 cells (Fig. 3e). Select DEGs were tested by qRT-PCR in WT/KO versus 6KR/KO MCF7 cells with or without Ex527 treatment. Expression of these genes was inhibited by Ex527 in WT/KO cells, but not 6KR/KO cells in which the expression was already at a higher level relative to the level in WT/KO cells (Fig. 3f). An analysis of select up-regulated DEGs by qRT-PCR using intron-targeting primers revealed that pre-mRNAs for these genes were also up-regulated (Supplementary Fig. 4d), in accordance with the notion that MED1-containing Mediator is involved in transcription, but not post-transcriptional RNA control. Next, DEGs of 6KR/KO versus WT/KO cells were compared with DEGs of *MED1* KO versus control cells by GSEA analyses. Using detected genes of *MED1* KO versus MED1 control as the rank, the GSEA revealed that the up-regulated genes in 6KR/KO cells were enriched in the side of down-regulated genes of *MED1* KO cells, while the down-regulated genes in 6KR/KO cells were enriched in the side of up-regulated genes of *MED1* KO cells (Fig. 3g). A GSEA analysis for 6KR/KO versus WT/KO non-differential genes showed no enrichment (NES = −0.795, p = 1.00). In parallel, using detected genes from 6KR/KO versus WT/KO as the rank, GSEA analysis for DEGs in *MED1* KO cells showed consistent results (Supplementary Fig. 4e). Here, GSEA analysis for *MED1* KO versus control non-differential genes showed no enrichment (NES = −0.948, p = 0.86). By correlating the fold-change of each individual gene from 6KR/KO versus WT/KO and from *MED1* KO versus *MED1* WT control tests, the inverse correlation was also established (Fig. 3h). These results lead us to envision that non-acetylatable 6KR MED1 acts as a more active form of MED1 for gene activation.

### The MED1 acetylation mutant affects genomic occupancy of Mediator and RNA Pol II in ER^+^ BC cells

In the next effort to investigate the mechanism by which non-acetylatable 6KR MED1 enhances MED1 target gene expression, we examined the genomic occupancies of 6KR and WT MED1 and their associated factors. The WT/KO and 6KR/KO cells used for ChIP-seq showed comparable levels of expression of ectopic (HA -tagged) WT and 6KR MED1, MED17, Pol II (RPB1, the largest subunit of Pol II) and ER, as well as efficient depletion of endogenous MED1 (Supplementary Fig. 5a). We performed ChIP-seq for HA, MED17 and Pol II in biological replicates in WT/KO and 6KR/KO cells respectively and verified that the replicates for each factor in each cell type were highly correlated (Supplementary Fig. 5b). MED1/MED17/Pol II (RPB1)-occupied peaks identified from the ChIP-seq data from untreated MCF7 cells (Fig. 1d, e) were used as pre-defined MED1/MED17/Pol II-occupied sites that were monitored in the following studies. Densities of ER, HA-WT MED1, HA-6KR MED1, MED17 and Pol II (RPB1) from ChIP-seq of 6KR/KO and WT/KO cells at the pre-defined MED1^+^ peaks were examined. An appreciable enhanced occupancy of HA-6KR MED1 relative to HA-WT MED1 was detected at MED1 and MED17 co-occupied peaks (regardless of Pol II occupancy) (Fig. 4a). Pol II (RPB1) also showed higher occupancy at the MED1^+^MED17^+^Pol II^+^ peaks in 6KR/KO cells relative to WT/KO MCF7 cells (Fig. 4a). In contrast, MED17 and ER occupancies showed little or no change at these peaks (Fig. 4a). By restricting the analyses to ER- and Mediator-occupied active enhancers (pre-defined as ER^+^ MED1^+^ MED17^+^ H3K27ac^+^ H3K4me^+^ H3K4me3^−^ peaks), we found analogous results for occupancies of the factors (Fig. 4b). This analysis was also performed for all active promoters (monitored by H3K4me3^+^ within ±1kb of the TSS) and similar changes in occupancies of these factors were observed and are shown in the metaplots (Fig. 4c) and heatmaps (Fig. 4d). Occupancy of these factors in regulatory regions of a representative gene (*TFF1*) is shown in IGV (Integrative Genomics Viewer) genomic snapshots (Fig. 4e).

**Fig. 4.**
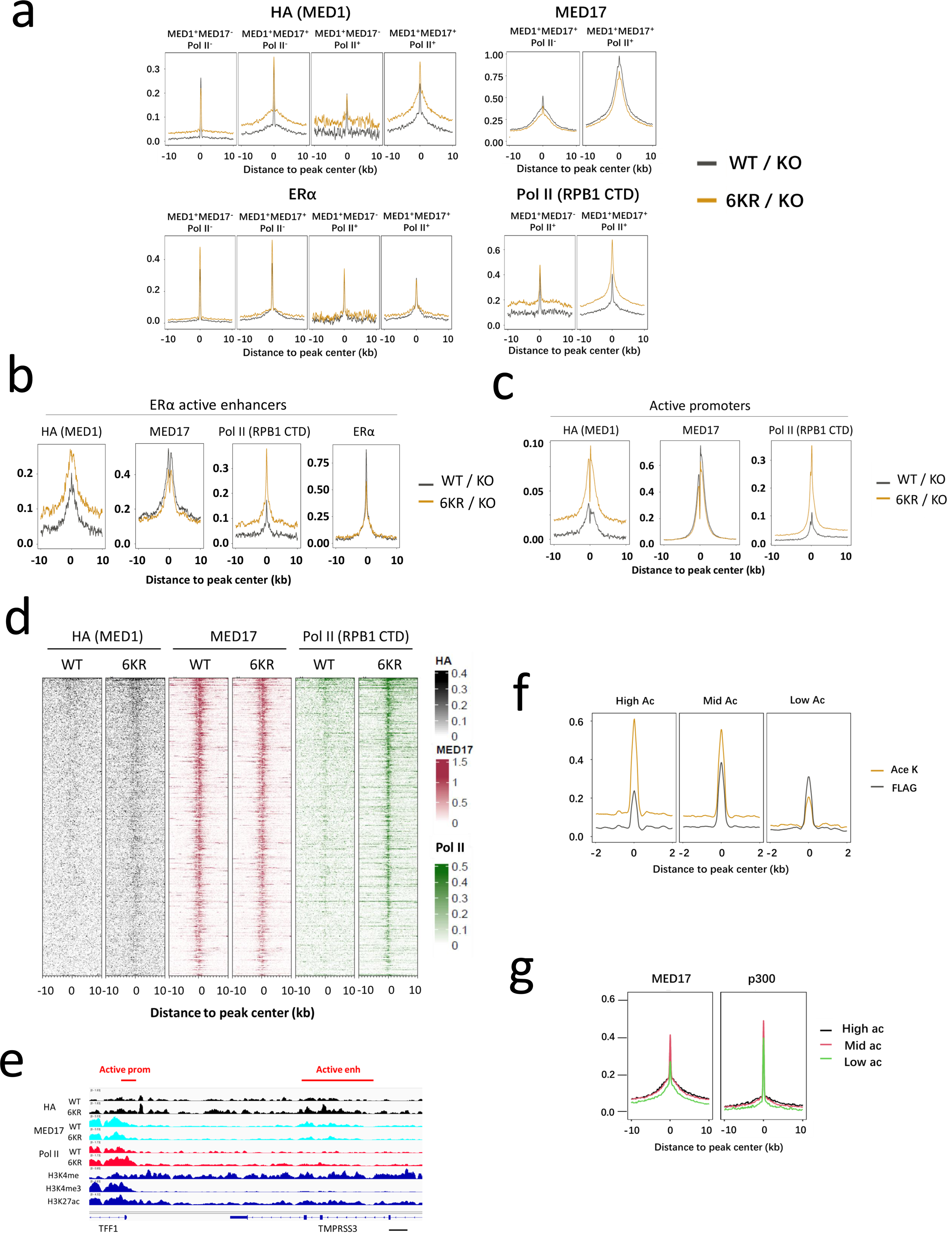
Chromatin occupancy of Mediator and RNA Pol II is altered by MED1 acetylation mutant and associated with endogenous acetylation level. (a, b) Density metaplots for pre-defined MED1 peaks (further classified according to their co-occupancy with MED17 and Pol II) (a) and ER^+^ active enhancers (b) showing occupancy of ERα, (ectopic) MED1, MED17, and Pol II with comparison between 6KR/KO and WT/KO cells. (c, d) Metaplots (c) and heatmaps (d) of active promoters showing occupancy of (ectopic) MED1, MED17, and Pol II with comparisons between 6KR/KO and WT/KO cells. (e) IGV snapshot of regulatory regions of a representative gene, *TFF1*, with occupancy of HA (MED1), MED17 and Pol II shown for WT/KO and 6KR/KO cells. Epigenetic marks to identify enhancers and promoters are also shown. (f) Metaplots of occupancy of FLAG and acetylation for pre-defined MED1 peaks in High Ac, Mid Ac and Low Ac groups. (g) Metaplots of occupancy of MED17 and p300 for pre-defined MED1 peaks with comparison among High Ac, Mid Ac and Low Ac groups.

Motivated by the differential genomic occupancies of 6KR and WT MED1, we next explored the genomic distribution of acetylated MED1 (relative to total MED1) in MCF7 cells. Limited by the availability of acetylation site-specific antibodies, we employed a sequential ChIP as an alternative. Thus, anti-FLAG immunoprecipitates from MCF7 cells expressing ectopic FLAG-tagged MED1 were eluted with FLAG peptide and followed with a second immunoprecipitation with pan-anti-acetylated-lysine antibodies (Supplementary Fig. 5c). This procedure enriches not only acetylated FLAG-MED1, but also other acetylated MED1 binding factors. However, the acetylation level of MED1 and/or its binding proteins at each of the pre-defined MED1 peaks (according to ChIP-seq of untreated MCF7 cells) can be evaluated by comparing the read densities from sequencing of 1^st^ and 2^nd^ ChIP samples. In accordance, we stratified the MED1 peaks by ranking the 2^nd^ ChIP/1^st^ ChIP ratios of read densities and taking the upper 75%, middle 50% and lower 25% percentiles, respectively (referred as High Ac, Mid Ac and Low Ac groups) (Fig. 4f, Supplementary Fig. 5d). The FLAG-MED1 density was lower for the High Ac group, which was accompanied by higher MED17 occupancy (Fig. 4f, g, Supplementary Fig. 5d). The p300 occupancy was lower for the Low Ac group but the difference was marginal (Fig. 4g, Supplementary Fig. 5d), raising the possibility that, in addition to p300, histone deacetylases may also play a critical role for levels of acetylation of MED1 and associated factors. Low Ac MED1 peaks exhibit less MED17 co-occupancy (total of MED17^+^Pol II^+^ and MED17^+^Pol II^−^) (Supplementary Fig. 5e), and these peaks are associated with less active enhancers (Supplementary Fig. 5f).

At face value (but see below), the above results suggest a possible dissociation of MED17 (most likely indicating dissociation of the entire Mediator core module in view of the integral position of MED17 in the Mediator structure^5,6^) and elevation of Pol II at active promoters in 6KR/KO cells or at low acetylated MED1 peaks in normal cells, being suggestive of a distinct composition or conformation of the Mediator/PIC at these loci. In this regard, it is likely that a conformational alteration of the Mediator/PIC is responsible for the reduced recognition by anti-MED17 antibodies as reflected in the ChIP-seq results. The up-regulation of MED1-dependent genes in MED1-deficient cells upon re-expression of the non-acetylatable 6KR mutant MED1 (Fig. 3g, h; Supplementary Fig. 4e) could be enabled by a novel MED17/Mediator dissociation or rearrangement mode of PIC function in transcription initiation and/or re-initiation.

### The MED1 acetylation mutant leads to sub-TAD chromatin unfolding

Gene transcription, associated with communication between cis-regulatory elements (promoters and enhancers), occurs within a dynamic 3D chromatin microenvironment^35^, but the functional association of transcription and 3D chromatin organization remains a matter of active debate^36^. Motivated by recent reports that acute depletion of MED14 or RPB1 significantly alters 3D chromatin interactions in human cells^13,14^, we hypothesized that MED1 acetylation is leveraged for the dynamic control of 3D chromatin organization. Employing the HiCAR method that uses Tn5 transposase and chromatin proximity ligation to capture open chromatin-involved interactions at 5 kb resolution^37^, we sequenced samples prepared from WT/KO and 6KR/KO MCF7 cells under normal growth conditions (Supplementary Fig. 6a).

The ATAC (Assay for Transposase-Accessible Chromatin) signal (generated by HiCAR) with 2 kb bins spanning the entire genome showed strong correlation between WT/KO and 6KR/KO cells (Supplementary Fig. 6b), indicating comparable chromatin accessibility. Compared with a published dataset of ATAC-seq for MCF7 cells (GEO Series GSE202511), the ATAC peaks from either WT/KO or 6KR/KO cells showed high overlap with published ATAC peaks (Supplementary Fig. 6b). In WT/KO and 6KR/KO MCF7 cells, 28,363 chromatin interactions were detected by HiCAR in total, and compared with published Hi-C study for MCF7 cells (GEO Series GSE99541), either WT/KO or 6KR/KO cells showed high overlap for chromatin interactions (Supplementary Fig. 6c). Published Hi-C results from MCF7 cells (GEO series GSE99541, no E2 control and E2-treated), MDA-MB-231 cells (ER^−^ BC cells, GEO Series GSE223785), and THP-1 cells (acute monocytic leukemia cells, GEO Series GSE226673) were collected and analyzed by the HiCRep method^38^ in order to compare the contact matrices of these datasets with our HiCAR results from WT/KO and 6KR/KO MCF7 cells (Fig. 5a). The interaction profiles of WT/KO and 6KR/KO MCF7 cells resembled one another, and both showed high correlation with published MCF7 results. These results contrast with the lower correlation of Hi-C results for MDA-MB-231 and THP-1 cells, demonstrating that the chromatin interactions identified by our HiCAR analyses faithfully reflected the biological feature of MCF7 ER^+^ BC cell line (Fig. 5a). A comparison of chromatin architecture of select regions is shown at various resolutions (Fig. 5b) where an alteration of interactions near a typical 6KR/KO DEG, *TFF1*, is observed (Fig. 5b).

**Fig. 5.**
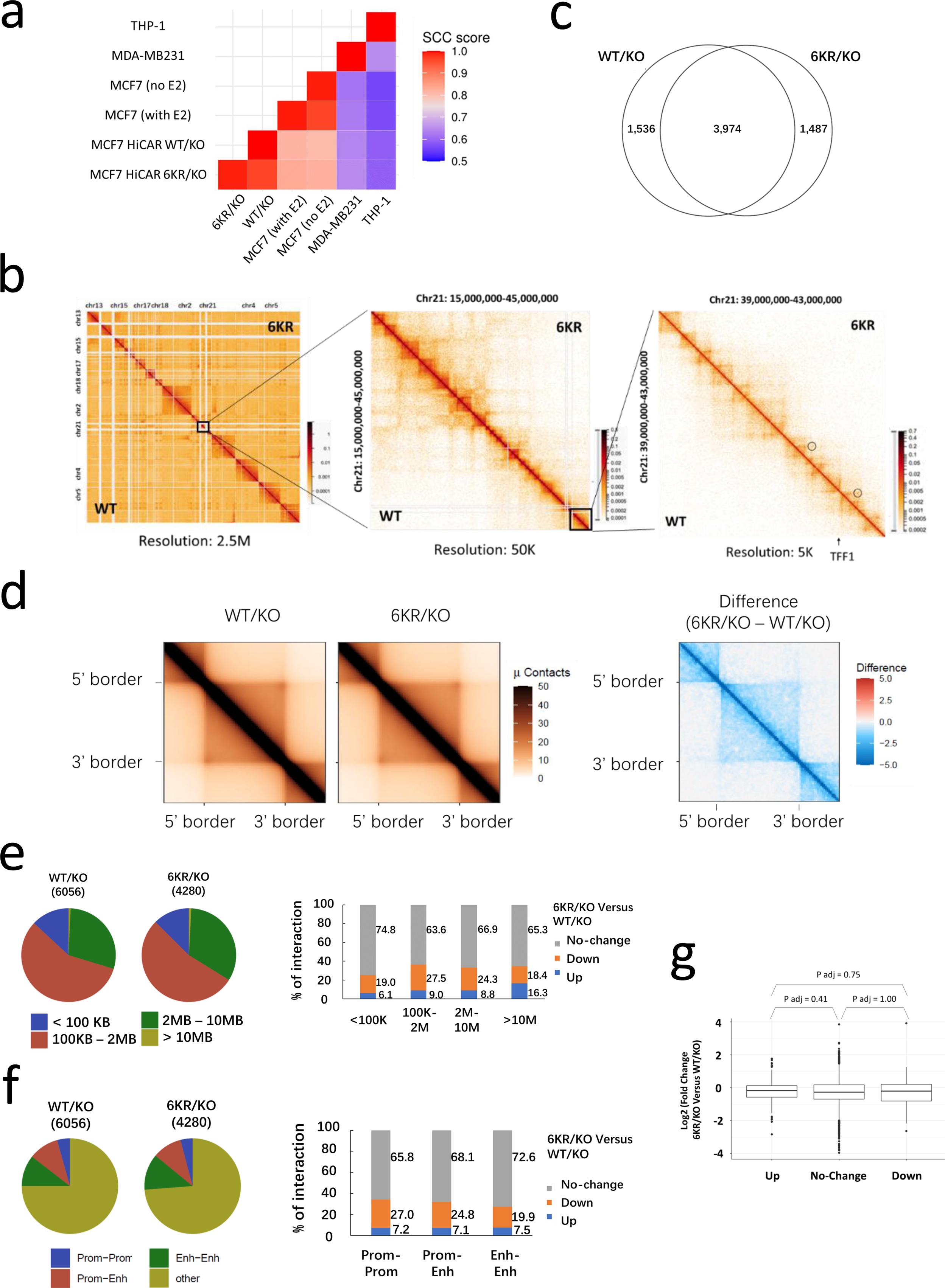
The non-acetylatable 6KR MED1 mutation alters sub-TAD chromatin interactions. (a) A heatmap of SCC (Stratum-adjusted Correlation Coefficients) scores of HiCRep analysis comparing chromatin contact matrices of HiCAR of WT/KO and 6KR/KO MCF7 cells and published Hi-C of MCF7, MDA-MB-231 and THP-1 cells. Published Hi-C data use: MCF7: GEO series GSE99541, taking union of 2 samples of GSM2645712 and GSM2645713 as no E2 control, taking union of 2 samples of GSM2645714 and GSM2645715 as E2-treated; MDA-MB-231: GEO Series GSE223785, taking union of 2 samples of GSM6995361 and GSM6995362; THP-1: GEO Series GSE226673, taking union of 2 samples of GSM7081346 and GSM7081347. (b) Heatmaps of chromatin interactions identified from WT/KO and 6KR/KO cells by HiCAR in a select region with 2.5 Mb, 50 kb and 5 kb resolutions. A typical ERα and MED1 target gene, *TFF1*, is pointed and differential interactions near the *TFF1* are indicated. (c) Venn diagram showing numbers of TADs identified from WT/KO and 6KR/KO MCF7 cells. (d) Aggregate TAD analysis (ATA) (left) with differential plot (right) showing comparable TADs identified from WT/KO and 6KR/KO MCF7 cells and global reduction of sub-TAD chromatin interactions. For (c) and (d), TADs were identified with a window size of 250 kb. (e) Size distribution of chromatin interactions with MED1 occupying at least one anchor in WT/KO and 6KR/KO MCF7 cells (left) and frequency of alteration (6KR/KO versus WT/KO) for each size category (right). (f) Constitution of MED1-positive chromatin interactions for their anchoring at active promoters (referred as Prom), active enhancers (referred as Enh) or other regions in WT/KO and 6KR/KO MCF7 cells (left) and frequency of alteration (6KR/KO versus WT/KO) for each category (right). For (e) and (f), 1.5-fold change was used as cut-off for altered chromatin interactions. (g) A box plot showing fold-change values of promoter-anchored chromatin interactions in 6KR/KO versus WT/KO cells with categorization according to the promoter association with DEGs from 6KR/KO versus WT/KO cells (p=0.05 as cut-off). P adj values between any two groups are shown. For all panels, chromatin interactions were identified in 10 kb bins as described in Methods.

Topologically associating domains (TADs) are 3D chromatin structures that are highly conserved across species and rarely altered in cells^39^. About 2,200 TADs can be detected in mammalian chromatin by routine Hi-C methods^39^, while about 5,500 TADs were called in each of our WT/KO and 6KR/KO samples. The average sizes of TADs observed in WT/KO and 6KR/KO samples were 531 kb and 536 kb, respectively, being close to the initial characterization of TAD size as 880 kb^39^. The TADs of either WT/KO or 6KR/KO cells showed high overlap with TADs identified from published Hi-C results for MCF7 cells (GEO Series GSE99541) (Supplementary Fig. 6d). The majority of TADs was called in both WT/KO and 6KR/KO samples (Fig. 5c for overlap of TAD boundaries). Comparison of the TADs^40^ revealed that 78.6% of those in WT/KO cells showed no change in 6KR/KO cells while 17.0% showed border-shift -- but the split or merging of TADs or the strength change of the overlapped TADs are all rare (Supplementary Fig. 6e). In contrast to the stable nature of TADs, a differential plot generated from aggregate TAD analysis (ATA) indicated that interactions inside TADs (including short-range interactions between regulatory elements of genes) are diminished in the 6KR/KO sample relative to the WT/KO sample (Fig. 5d).

In either WT/KO or 6KR/KO MCF7 cells, the vast majority of observed chromatin interactions are 100 kb to 2 Mb length, while a very limited number of interactions extend over 10 Mb (Supplementary Fig. 6f). In contrast, the interactions with at least one MED1-occupied anchor show a higher distribution in the 2 Mb to 10 Mb range, the typical TAD size, while the proportion of interactions in the 100 kb to 2 Mb range is reduced (Fig. 5e). Of all the chromatin interactions detected, only a limited proportion is anchored at active promoter or enhancer sites (Supplementary Fig. 6g). However, there is an increased proportion of MED1-occupied interactions at these cis-regulatory elements (Fig. 5f), indicating a closer association with gene regulation. For MED1-occupied interactions in most categories described here (length categories or anchor categories), there is an analogous trend of down-regulation in 6KR/KO relative to WT/KO cells. This is evidenced by detection of a significant proportion of weakened interactions and only rare strengthened interactions (although more than half of the interactions show no difference) (Fig. 5e, f). These results are consistent with the widespread suppression of sub-TAD chromatin interactions (Fig. 5d).

To gain insight into the specific association of 3D chromatin interactions with gene regulation under the control of the MED1 6KR mutation, we next divided MED1-occupied chromatin interactions into three groups according to their association with up-regulated, down-regulated or unhanged genes in 6KR/KO versus WT/KO MCF7 cells. Aggregate peak analysis (APA) showed down-regulation of all three groups of interactions in 6KR/KO versus WT/KO cells (Supplementary Fig. 6h, comparing either the Z score or the APA value that is shown in each panel). In light of the importance of interactions between promoters and distal enhancers in gene regulation^35^, we analyzed only those interactions anchored at regions containing gene promoters and divided them similarly into three groups according to their association with up-regulated, down-regulated or unchanged genes in 6KR/KO versus WT/KO cells. The promoter-anchored interactions from any group showed no observable differences from the other two groups for the trend of down-regulation in 6KR/KO versus WT/KO cells (comparing fold-change values) (Fig. 5g). These findings bring into question the current, relatively straightforward hypothesis that MED1 acetylation (or 6KR mutation) controls activation of genes by re-structuring their surrounding 3D architecture (but see discussion for the limitation of our study.)

The sequential ChIP-seq experiment (Fig. 4f, g; Supplementary Fig. 5c-f) divides MED1-occupied sites according to the acetylation level of FLAG-MED1 or a potential binding factor. Considering the anchoring sites of chromatin interactions identified by HiCAR, a much higher proportion of the interactions is anchored at High/Mid Ac MED1 peaks than at Low Ac MED1 peaks (Supplementary Fig. 6i). This result supports the observation of reduced sub-TAD chromatin interactions in 6KR/KO (non-acetylatable) cells relative to WT/KO cells and indicates that HiCAR-identified open chromatin interactions are more abundant at the sites with higher acetylation of MED1 or its associated transcription (co)factors.

### IDR acetylation determines the phase behavior of MED1 as well as Pol II sequestration

As the 6 major MED1 acetylation sites are located in the MED1 IDR, and as accumulating evidence supports critical roles for IDRs in LLPS and condensation of proteins^41^, we anticipated that acetylation would change the physio-chemical properties of the MED1 IDR (i.e., a transition of phase behavior) and thus impact MED1 condensation. To address this issue, a HIS- and mEGFP-fused MED1 IDR segment (residues 963-1320 predicted by VLXT, Supplementary Fig. 3c) with either 6KR or 6KQ (lysines replaced by glutamines) mutations was expressed in *E.coli* and purified with Ni-NTA agarose beads (Supplementary Fig. 7a).

Following imidazole removal by dialysis, 6KR and 6KQ IDRs both formed droplets at 40 μM in 100 mM NaCl and 10% polyethylene glycol (PEG, non-ionic crowding agent) -- with 6KR droplets being larger than 6KQ droplets (Fig. 6a). However, turbidity measured as absorbance at 600 nm demonstrated that 6KQ IDR is more prone to LLPS when protein or PEG concentrations are lower (Supplementary Fig. 7b, c). Fluorescence recovery after photobleaching (FRAP) showed faster recovery of 6KQ IDR relative to 6KR IDR (Fig. 6b, c), further supporting the idea that both the acetylation-mimicking 6KQ IDR and the non-acetylatable (or deacetylation-mimicking) 6KR IDR undergo condensation – but with the 6KQ IDR exhibiting more fluidity (liquid-like feature) and the 6KR IDR exhibiting more solid/gel-like features. The distinct physio-chemical nature of the 6KR and the 6KQ IDR for LLPS was also tested in a co-condensation assay. The mCherry-fused 6KR and 6KQ MED1 IDRs were purified from *E.coli* (Supplementary Fig. 7a) and their co-condensations with mEGFP-fused 6KR and 6KQ MED1 IDRs were imaged. Although the co-condensates of mEGFP-6KR + mCherry-6KR and mEGFP-6KQ + mCherry-6KQ, respectively, showed homogenous mixing, the co-condensates of mEGFP-6KR + mCherry-6KQ and mEGFP-6KQ + mCherry-6KR showed a heterogeneity that was observed as a separation of mEGFP- and mCherry-dominated regions and supported by pixel-based computation (Fig. 6d).

**Fig. 6.**
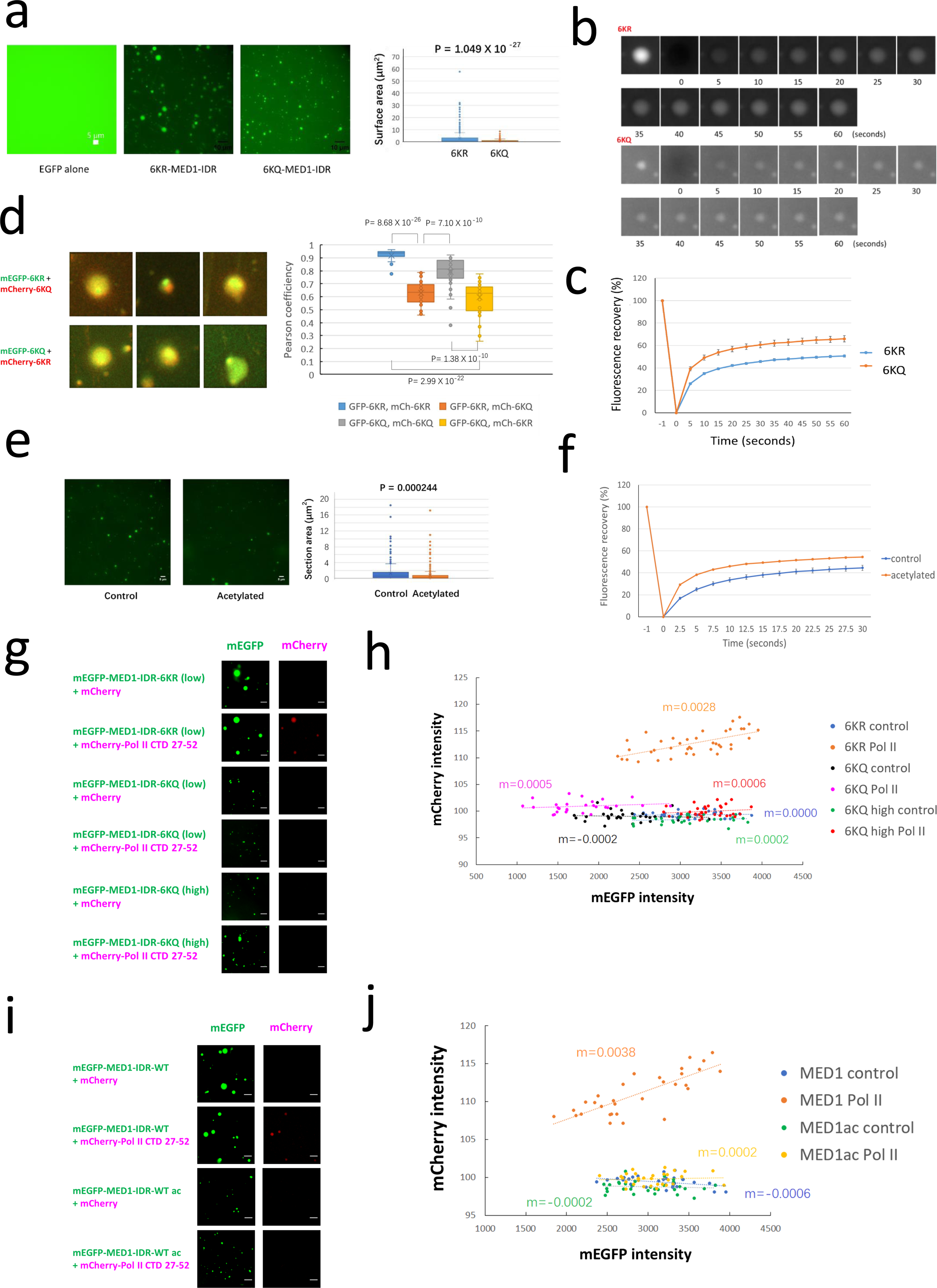
Acetylation alters physio-chemical properties of the MED1 IDR for LLPS. (a) Larger droplets formed by the 6KR MED1 IDR compared to the 6KQ MED1 IDR (fused with mEGFP) at 40 μM in a buffer containing 100 mM NaCl and 10% PEG. Representative images (left) and box plots (right) of droplet size (surface area computed with 3D images) are shown. P value of t test is shown. (b, c) Faster recovery of 6KQ relative to 6KR IDR droplets observed by FRAP. Images for representative droplets (b) and recovery curves (c, data points report mean ± SE) are shown. 6KR and 6KQ IDRs were tested for 6 and 7 droplets, respectively. (d) Co-condensates of the 6KR/Q MED1 IDR (fused to mEGFP/mCherry) showing heterogeneous mixing of 6KR + 6KQ IDRs compared with 6KR + 6KR and 6KQ + 6KQ IDRs. Images of representative droplets (left) and pixel-based computations for Pearson co-efficiency with P values of t test (right) are shown. (e) Smaller droplets formed by CBP-treated compared to control WT MED1 IDR (fused with mEGFP) at 13.8 μM in a buffer containing 100 mM NaCl and 10% PEG. Representative images (left) and box plots for section area of droplets with P value of t test (right) are shown. (f) Faster recovery of droplets formed by CBP-treated versus control WT MED1 IDR revealed by FRAP. Data points report mean ± SE. Control and CBP-treated MED1 IDRs were tested for 6 droplets for each. (g-j) Sequestration of an mCherry-fused Pol II CTD heptad fragment (but not mCherry alone control) at 0.19 μM into condensates formed by mEGFP-fused 6KR (36 μM), but not 6KQ (36 or 76.5 μM) MED1 IDR (g, h), and mEGFP-fused control but not CBP-acetylated WT MED1 IDR (36 μM for each) (i, j). Representative images (g, i) and signal intensity for mCherry plotted as a function of signal intensity for mEGFP for each droplet (h, j) are shown.

Apart from testing 6KR and 6KQ MED1 IDRs, we also examined the extent to which the difference between 6KR and 6KQ MED1 IDRs reflects the effect of MED1 IDR acetylation. In this regard, WT MED1 IDR was purified from *E.coli* as previously, except that during purification the bead-bound MED1 IDR was subjected to acetylation by incubation with purified mouse CBP HAT domain and acetyl-CoA (Supplementary Fig. 7d, e). With elevated acetylation confirmed (Supplementary Fig. 7e), the CBP-treated WT MED1 IDR formed larger droplets (Fig. 6e) and exhibited a slower FRAP recovery rate (Fig. 6f, Supplementary Fig. 7f) -- revealing a parallel to the differential phase behavior between 6KR and 6KQ MED1 IDRs, and again indicating the effects of acetylation on LLPS. As expected, the WT MED1 IDR without enforced acetylation and the 6KR MED1 IDR behaved similarly in LLPS, as indicated by the recovery kinetics in FRAP assays (comparing Fig. 6c and 6f).

RNA Pol II and MED1 co-condensates at active transcription sites have been reported^42^ and Pol II enrichment in Mediator loci in cells is dependent on MED14^10^. To gain further insight, we purified a human Pol II CTD heptad fragment^43^ fused to mCherry from *E.coli* (Supplementary Fig. 7g) and imaged its co-condensation with mEGFP-MED1 IDR (Fig. 6g-j). At a concentration much lower than the MED1 IDR concentration, the Pol II CTD did not form homotypic condensates, but was enriched within 6KR MED1 IDR droplets. In contrast, we did not observe Pol II CTD enrichment in 6KQ MED1 IDR droplets, even at a higher concentration that supports formation of 6KQ MED1 IDR droplets comparable in size to 6KR MED1 IDR droplets (Fig. 6g). As indicated by the intensity plot, enrichment of the Pol II CTD positively correlated with 6KR MED1 IDR droplets (Fig. 6h). We also examined the enrichment of Pol II CTD heptads within droplets of WT MED1 IDR with or without CBP acetylation (Supplementary Fig. 7d, e). At a low concentration, the Pol II CTD was only concentrated in droplets formed by the MED1 IDR without CBP acetylation (Fig. 6i). The positive correlation between Pol II CTD and non-acetylated MED1 IDR in droplets was supported by an intensity plot (Fig. 6j). These in vitro studies establish that IDR acetylation significantly alters the phase behavior of MED1, which may potentially regulate target gene transcription through MED1 condensation and enrichment of the transcription machinery including Pol II.

## Discussion

As an essential transcription coactivator, the Mediator acts together with gene-specific transcription activators and the RNA Pol II machinery and other factors involved in various steps of transcription and mRNA processing to effect gene activation^1,8^. The Mediator may also work in concert with cohesin in establishing 3D chromatin architecture favorable for transcription in mammalian cells^13,44^ and yeast^16^. Therefore, the Mediator is posited at the interface between linear and spatial chromatin functions to drive gene activation. While not generally required for cell viability^45^, the ubiquitously expressed MED1 Mediator subunit is frequently monitored as a surrogate for Mediator and has been reported to have an essential role in transcription in certain cell types that include ER^+^ BC cells^3,46–48^. Consistent with the observation that histone modifying factors also functionally modify many transcriptional (co)activators^49^, we report that MED1 is acetylated with 6 major acetylation sites mapped to the IDR. Replacement of the endogenous MED1 by the non-acetylatable 6KR MED1 mutant in MCF7 cells accelerates cell growth and colony formation, and enhances the function of MED1 in gene activation. Our observations suggest that the 6KR mutation might exploit two mechanisms to achieve the enhanced MED1 target gene activation: (1) reorganization of Mediato and/or PIC complexes. This possibility is suggested by the observation that compared to WT MED1 cells, 6KR mutant cells show higher MED1 6KR and Pol II and lower MED17/Mediator occupancies, but unaltered ER occupancies, at active enhancers and promoters; (2) function of MED1 acetylation as a hub to coordinate Mediator-Pol II (PIC) assembly and sub-TAD chromatin folding to modulate gene expression (see the model in Supplementary Fig. 8). This possibility is suggested by our observation that despite the apparent lack of a specific association of 3D chromatin interactions with gene regulation in MED1 6KR cells, broad inhibitory effects of the MED1 6KR mutant on sub-TAD chromatin interactions were observed by HiCAR.

The functional association of transcription and 3D chromatin organization is still fervently debated. It has been reported that transcription inhibitors exert little effect on 3D chromatin architecture^12^ and, further, that acute depletion of the MED14 scaffolding subunit of Mediator^9,10^, a subunit of general transcription factor TFIID^50^, or a Pol II subunit^11^ remained largely inconsequential for overall chromatin architecture and most enhancer-promoter interactions. However, recent studies employing a single-nucleosome resolution micro-C strategy revealed selective loss of enhancer-promoter interactions and rewiring of CTCF-dependent loops by acute depletion of MED14^13^ or RPB1^14^. In agreement with these findings of the critical roles of Mediator and Pol II, our HiCAR analyses demonstrated a widespread down-regulation of sub-TAD MED1-occupied chromatin interactions in MED1 6KR cells -- in particular, those anchored at active enhancers and promoters and spanning distances within the TAD scale (Fig. 5e, f). Interestingly, MED1-occupied chromatin interactions spanning over 10 Mb exhibited comparable up-regulation and down-regulation (i.e., un-changed in general) (Fig. 5e), supporting the notion that these ultra-long-range, multi-TAD-spanning chromatin interactions are regulated by a distinct mechanism^51,52^. In line with reports that active transcription is associated with a less compacted chromatin architecture in yeast^16^ and that Mediator (through its kinase module) helps to maintain less compacted chromatin architecture favorable for transcription^15^, our results suggest a transcription-permissive 3D chromatin microenvironment through which 6KR MED1 promotes gene activation. However, it awaits further investigation as to whether the decline of sub-TAD chromatin interactions in 6KR MED1-expressing cells reflects a specific cell cycle phase and/or alteration of chromatin compartmentalization. In our analysis, the inability to establish a specific association of 3D chromatin interactions with gene regulation under the control of MED1 6KR might be caused by the resolution limit of HiCAR (similar to Hi-C). A more convincing proof of that association will require complete identification of finer-scale chromatin interactions with cis-regulatory elements, which could be achieved through a more powerful micro-C based strategy.

The involvement of MED1 IDR acetylation in gene regulation through PIC-related and 3D chromatin mechanisms might be manifested, at least in part, through LLPS. MED1 forms biomolecular condensates that enrich MED1 along with other transcription machinery components, including Pol II, to facilitate gene activation^42^. The 6KR (or control WT) MED1 IDR exhibits LLPS properties distinct from those of the 6KQ (or CBP-acetylated WT) MED1 IDR -- including the size and fluidity of droplets and heterogeneity within co-condensates. The observation of Pol II CTD sequestration specific to 6KR and WT MED1 IDR droplets is further suggestive of the distinct phase behavior of the MED1 IDR (with or without acetylation) in gene regulation. Previously, a MED14-dependent sequestration of Pol II into large Mediator foci (possibly the condensates) in cells was observed and, interestingly, the enriched Pol II was shown to be hypo-phosphorylated (not fully activated)^10^. Although acetylation has been reported to affect the phase behavior of various cytosolic and nuclear proteins^53,54^, the present data provide the first example of a transcription machinery component whose condensation is modulated by acetylation. Lysine acetylation in the MED1 IDR could reduce polarity and increase the contribution of hydrophobic interaction in condensates, which could potentially result in a transition to a more solid/gel-like status and less incorporation of Pol II. This molecular grammar of IDR may serve as a novel mechanism of MED1 action in Mediator, in addition to the dynamic MED1-MED4-MED9 Rod-Elbow tether revealed by recent cryo-EM studies^5,6^.

Overall, this study has identified a pleiotropic potential of MED1 acetylation in controlling distinct linear and 3D gene regulation processes in ER^+^ BC cells. Our findings are highly instrumental in revealing how acetylation governs the function of a transcriptional cofactor through transition of its phase behavior in orchestrating assembly of the transcription machinery and chromatin architecture. Moreover, as a coordination hub for integrating multiple critical mechanisms, MED1 acetylation may offer a distinctive therapeutic window for treating diseases, such as ER^+^ breast cancer, with MED1/Mediator-related transcription control.

## Acknowledgements

The authors thank members of the Roeder Lab for technical assistance, helpful discussions, and comments regarding the manuscript. We thank Dr. Henrik Molina (Director) and colleagues in The Rockefeller University Proteomics Resource Center for assistance with protein mass spectrometry analyses; Drs. Priyam Banerjee, Christina Pyrgaki and Alison North (Director) in The Rockefeller University Bio-Imaging Resource Center for assistance with droplet imaging and FRAP assays; Dr. Thomas Carroll (Director) and colleagues in The Rockefeller University Bioinformatics Resource Center for assistance with sequencing data analysis; and Dr. Rui Gong in Dr. Gregory Alushin Lab at The Rockefeller University for assistance with bacterial cell lysis by high-pressure homogenization; and Dr. Joe Rodriguez in Dr. Hermann Steller Lab at The Rockefeller University for assistance with using a microplate reader. We also thank Drs. Xiaolin Wei and Yarui Diao at Duke University for guidance on HiCAR experiments; Dr. Yanshan Fang at Shanghai Institute of Organic Chemistry, Chinese Academy of Sciences for advice on LLPS assays; and Dr. Fajun Yang at Albert Einstein College of Medicine for sharing MED15 plasmids. This work was supported by NIH Grants CA234575 and AI148387 to R.G.R.

---

**Supplementary Fig. 1.**
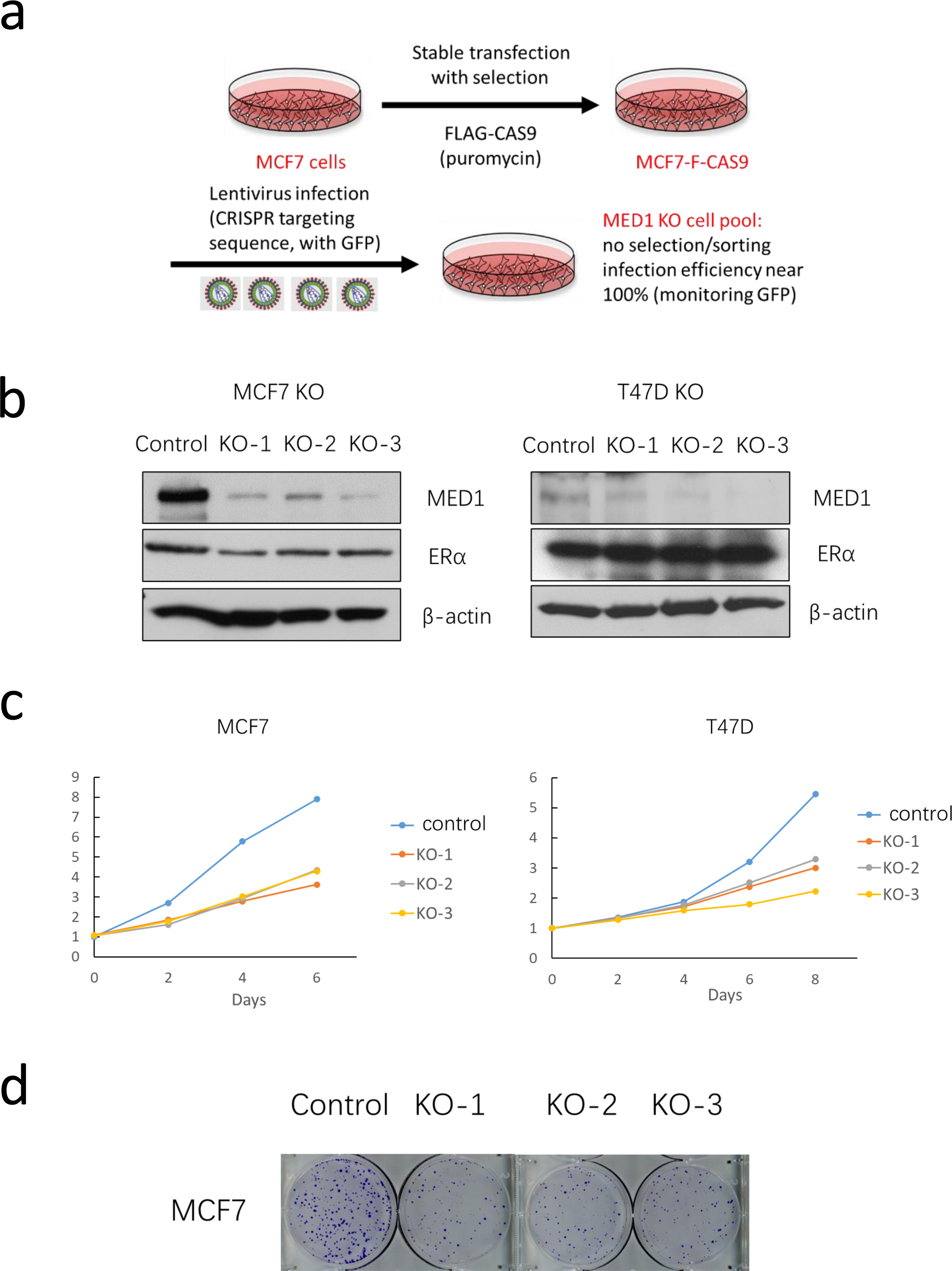
MED1 is essential for growth of ER^+^ breast cancer cells. (a) Workflow depicting the depletion of MED1 by CRISPR/Cas9. Cas9 (FLAG-tagged) was stably expressed in cells and lentiviral infection was used to introduce plasmids expressing gRNAs targeting *MED1* exons into pools of cells to deplete MED1. (b) Immunoblotting confirming depletion of MED1 by targeting 3 independent regions (KO-1, 2, and 3) in MCF7 (left) and T47D (right) cells, as well as comparable ERα levels in control and KO cells. (c) Cell growth attenuated by MED1 depletion in either MCF7 (left) or T47D (right) cells, measured by MTT assay. (d) Attenuated colony formation in MCF7 cells with *MED1* KO compared with control, observed by staining with crystal violet for cells growing for 16 days after initial seeding.

**Supplementary Fig. 2.**
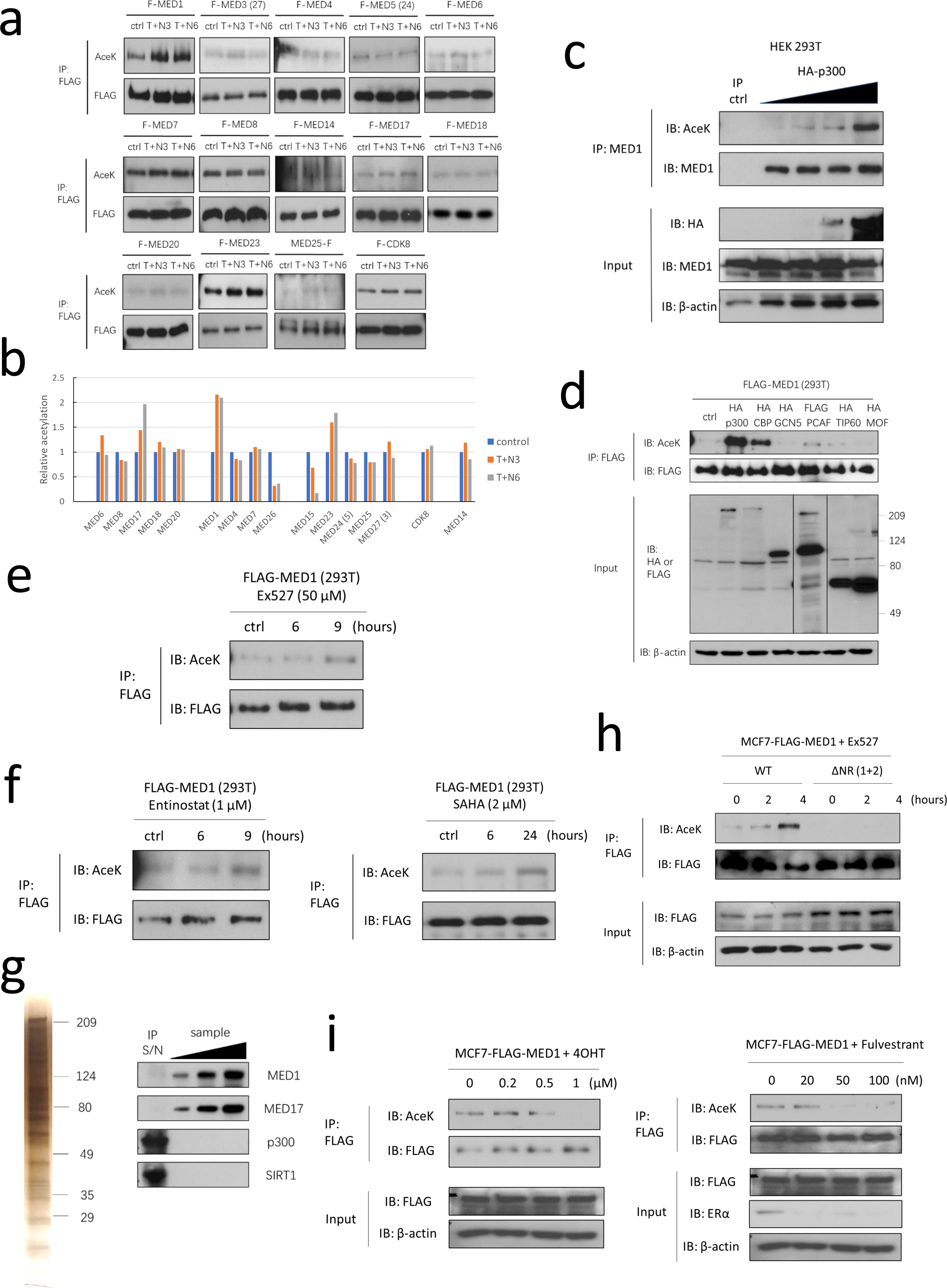
MED1 acetylation is controlled by p300/CBP histone acetyltransferases, histone deacetylases, and ER ligands. (a, b) Detection of acetylation of FLAG-tagged Mediator subunits transiently expressed in HEK293T cells treated with Trichostatin A (TSA, 0.5 μM for 24 hours) and nicotinamide (NAM, 0.5 mM for 3/6 hours), referred to as T+N3 or T+N6 in the panel. Immunoblotting images (a) and densitometry quantification with AceK immunoblot signals normalized by the FLAG signals (b) are shown. (c) Enhanced acetylation of endogenous MED1 from HEK293T cells with overexpression of p300, where the effects of variable p300 concentrations were tested. IgG was used as an IP control. (d) Enhanced acetylation of ectopically expressed FLAG-MED1 following overexpression of p300 (human) or CBP (mouse), but not GCN5, PCAF, TIP60 and MOF (all human), in HEK293T cells. (e, f) Enhanced acetylation of ectopically expressed FLAG-MED1 in HEK293T cells treated with Ex527 (e), Entinostat or SAHA (f). (g) Silver staining (left) and immunoblotting (right) confirming the yield and integrity of Mediator purified from HeLa-S cells and the absence of p300 and SIRT1. (h) Enhanced acetylation of ectopically expressed WT FLAG-MED1 but not mutant with deletion of the 2 NR boxes (referred as delNR(1+2)) in MCF7 cells treated with Ex527 (100 μM). (i) Reduced acetylation of ectopically expressed FLAG-MED1 in MCF7 cells treated with 4OHT (left) or Fulvestrant (right) for 8 hours at concentrations indicated in the panels. For all panels except (b) and (g), acetylation of immunoprecipitated FLAG-MED1 or endogenous MED1 was monitored by immunoblotting with pan-anti-acetyl-lysine antibody as described in Methods.

**Supplementary Fig. 3.**
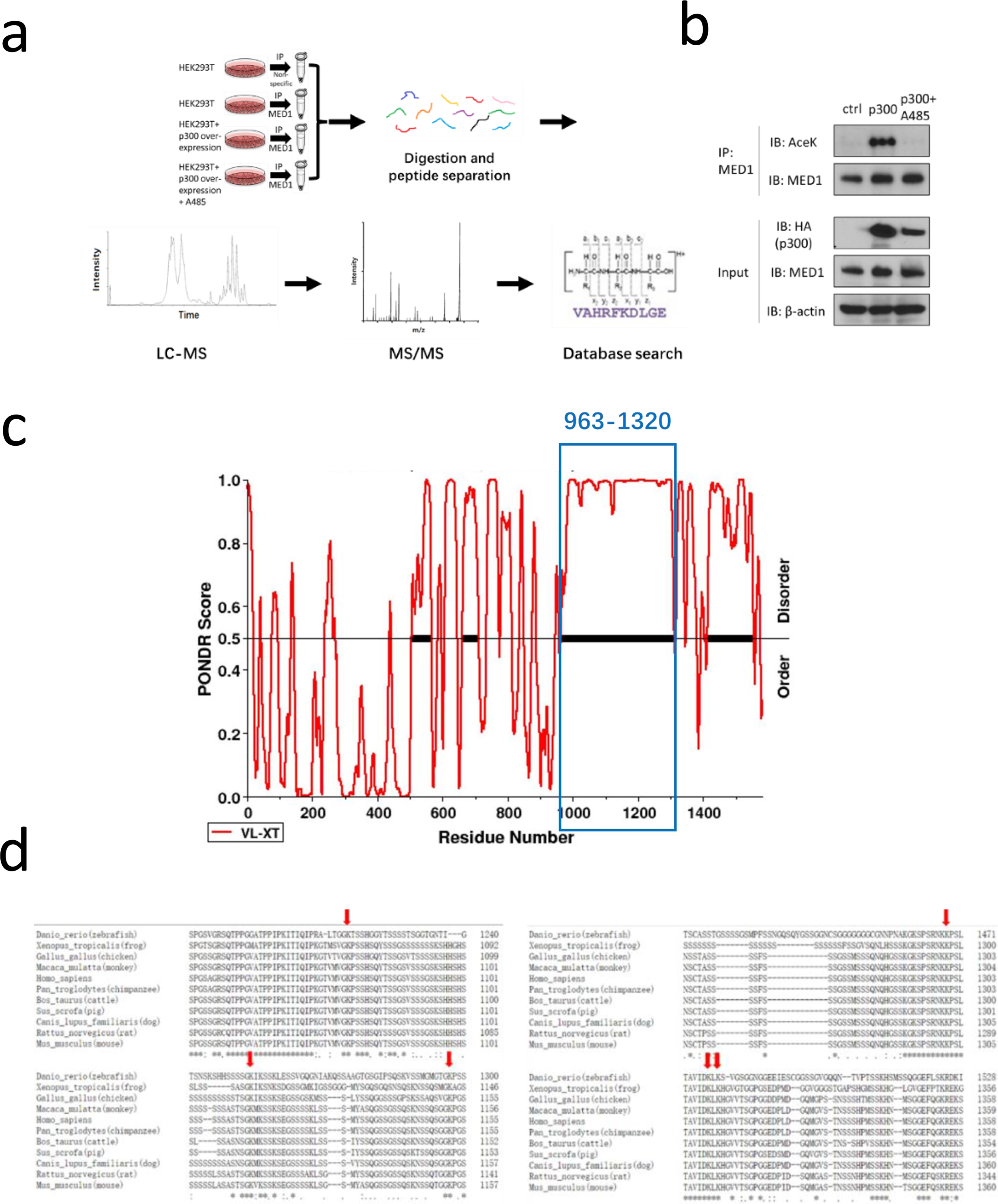
Identification of acetylation sites in the MED1 IDR. (a) Workflow for identification of acetylation sites from immunoprecipitated endogenous MED1 from HEK293T cells with overexpression of p300 with or without A485. (b) Immunoblotting confirming the overexpression of p300 and acetylation of MED1 in IP-Mass Spectrometry samples. (c) Analysis of MED1 protein sequence by VL-XT algorithm showing a long disordered region (963-1320 aa) with high PONDR score. (d) Multiple sequence alignments indicating that the 6 acetylation sites (arrows) in the MED1 IDR are conserved among species.

**Supplementary Fig. 4.**
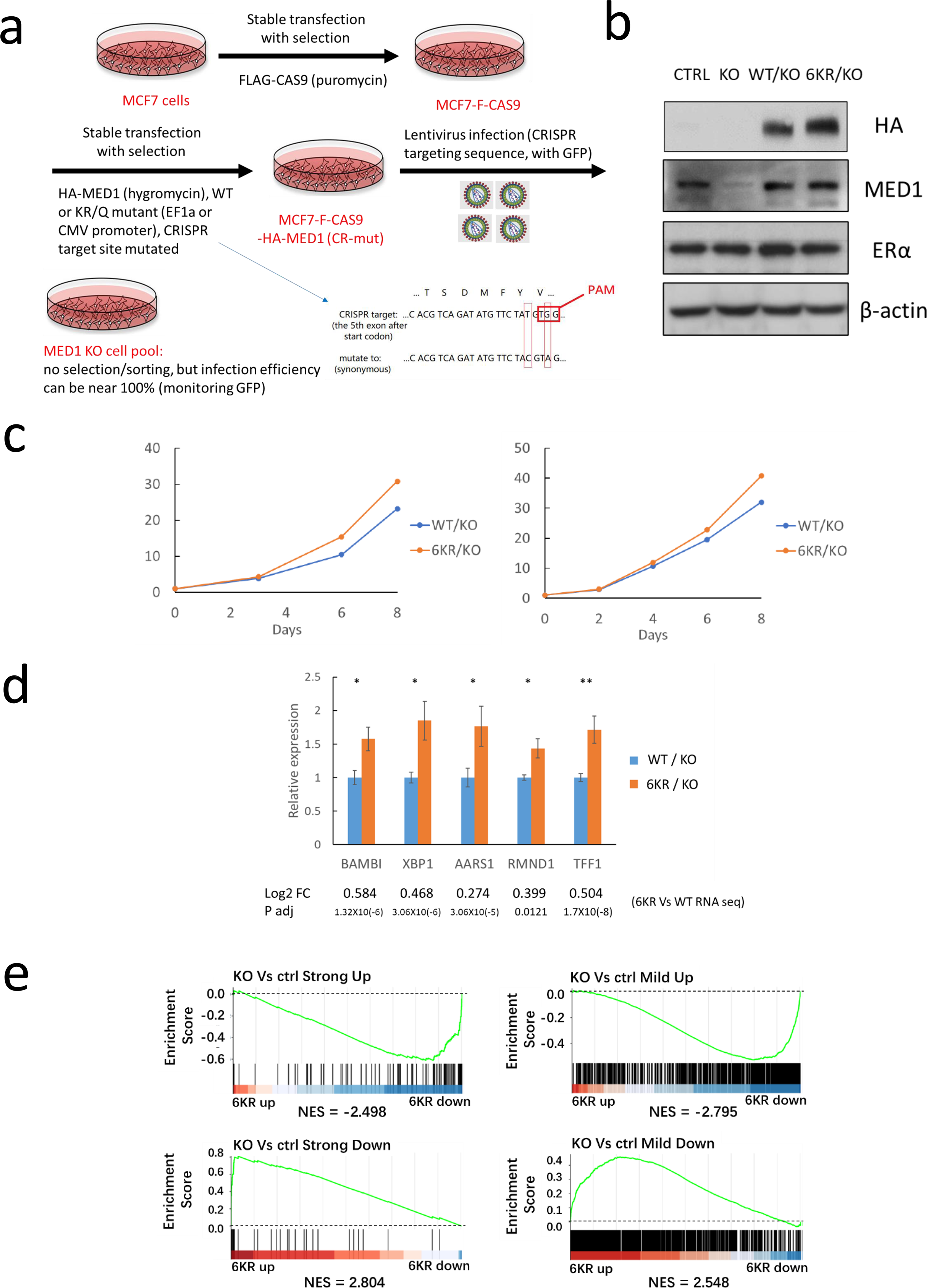
Replacing WT by 6KR MED1 enhances cell growth and MED1 target gene expression in MCF7 cells. (a) Workflow depicting the methods employed to replace endogenous MED1 with ectopically expressed WT or mutant MED1 in MCF7 cells. Lentiviruses expressing gRNAs targeting *MED1* were transfected into pools of cells expressing ectopic HA-tagged MED1 (WT or 6KR mutant) with CRISPR-mutated target sequences. (b) Immunoblotting confirming efficient depletion of MED1, expression of HA-tagged WT and 6KR MED1 (after CRISPR KO) at levels comparable to that of endogenous MED1, and no alterations in ERα abundance. (c) Two repetitions of the Fig. 3a analysis showing accelerated growth (monitored by CellTiter Blue assay) of 6KR/KO cells compared with WT/KO MCF7 cells. (d) Enhanced pre-mRNA expression of select up-regulated DEGs of 6KR/KO versus WT/KO cells tested by qRT-PCR with intron-targeting primers. The values for differential expression (fold-change and P adjusted) are also shown. N=6 for each group. Bars report mean ± SE. * p < 0.05, ** p < 0.01 (t test). (e) GSEA analyses revealing that the up-regulated (upper) and down-regulated (lower) genes in *MED1* KO versus control cells (strong or mild) are enriched, respectively, in the sets of down-regulated and up-regulated genes of 6KR/KO versus WT/KO cells.

**Supplementary Fig. 5.**
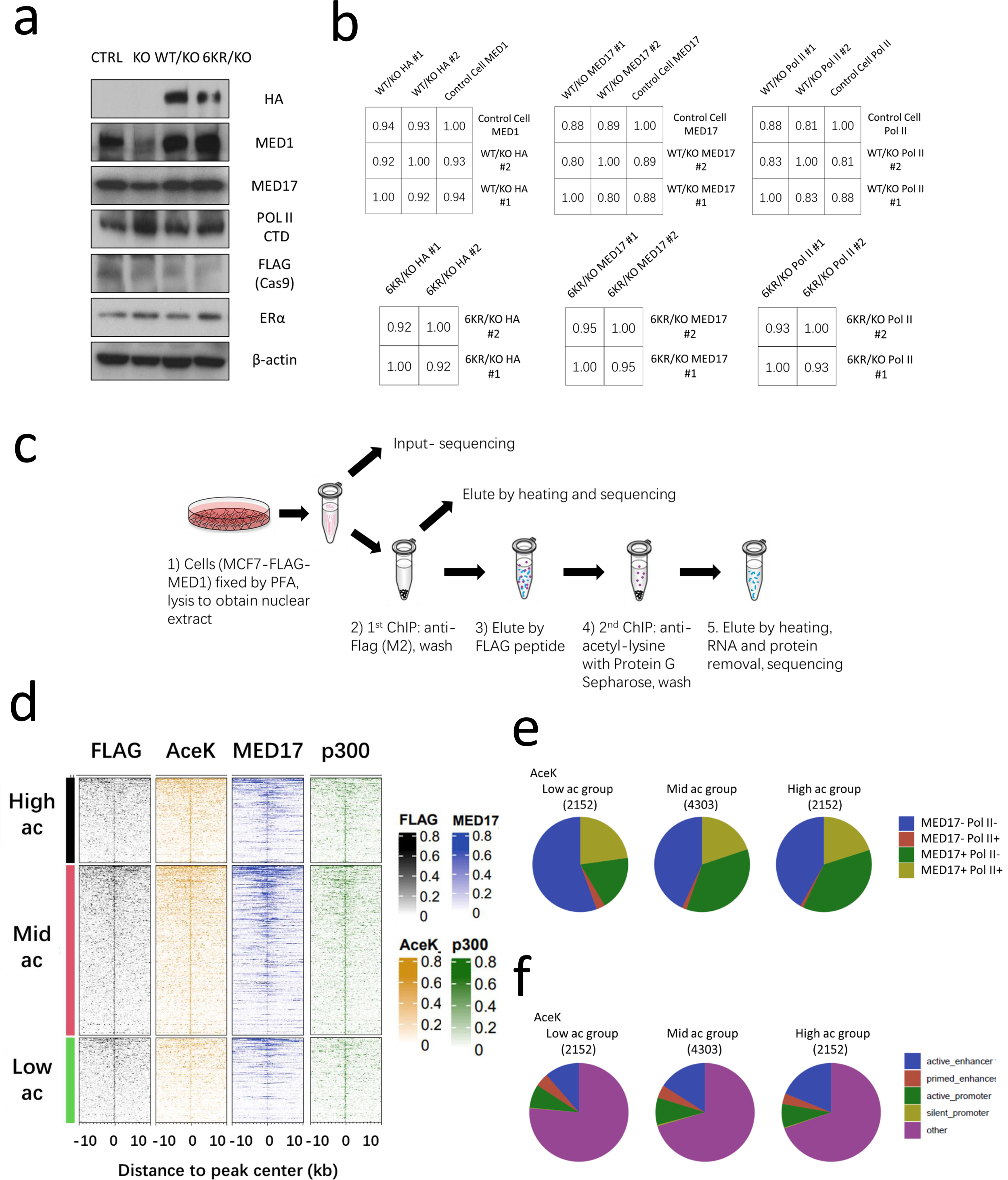
MED1 acetylation controls chromatin occupancy of Mediator and RNA Pol II. (a) Immunoblotting confirming depletion of endogenous MED1, equivalent expression of HA tag and other factors tested by ChIP-seq in WT/KO and 6KR/KO cells. (b) Results of Pearson correlation analysis at 5kb bins for ChIP-seq experiments of HA/MED1, MED17, and Pol II in replicate samples of WT/KO and 6KR/KO cells, and untreated MCF7 control cells. (c) Workflow depicting sequential ChIP-seq by anti-FLAG antibodies and pan-anti-acetyl-lysine antibodies to detect acetylation levels of FLAG-tagged MED1 (and its associated proteins) in genomic locations. (d) Range-based heatmap of the occupancy of FLAG-MED1, acetyl-lysine (2^nd^ ChIP), MED17, and p300 at pre-defined MED1 peaks that are stratified into 3 tiers based on relative acetyl-lysine density (lower 25%, middle 50% and upper 75% percentile, referred as Low Ac, Mid Ac, High Ac). (e, f) Pie diagram showing the constitution of High/Mid/Low Ac groups of MED1 peaks for occupancy of MED17 and Pol II (e), and types of regulatory elements (f).

**Supplementary Fig 6.**
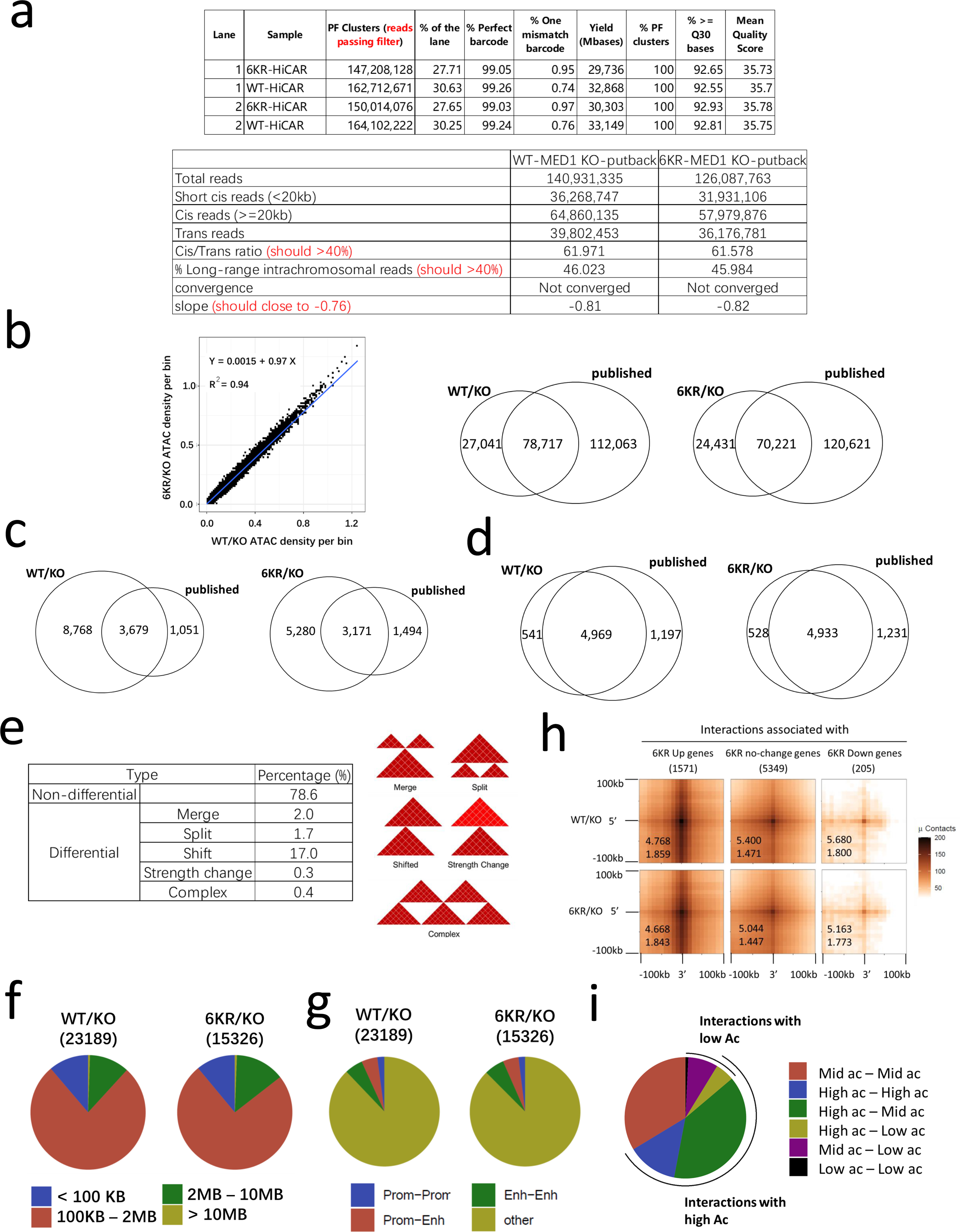
HiCAR analysis revealing alterations in 3D chromatin organization with MED1 acetylation mutation. (a) Yield of HiCAR sequencing experiments and the primary data analysis, including quality control. (b) Scatter plot showing strong correlation of ATAC signals with 2 kb bins spanning the entire genome from HiCAR results of WT/KO and 6KR/KO MCF7 cells (left). Venn diagram showing overlap of ATAC peaks from HiCAR results of WT/KO or 6KR/KO MCF7 cells with published MCF7 ATAC peaks (GEO Series GSE202511, Sample GSM6123339) (right). (c) Venn diagram showing overlap of chromatin interactions identified by HiCAR in WT/KO or 6KR/KO MCF7 cells with published MCF7 chromatin interactions. (d) Venn diagram showing overlap of TADs identified from HiCAR results of WT/KO or 6KR/KO MCF7 cells with TADs called from published MCF7 Hi-C data. TADs were identified with the window size of 250 kb. For comparison, a 50 kb shift for TAD boundaries was allowed. For (c) and (d), published MCF7 data (GEO Series GSE99541) was used by taking union from four experiments (GSM2645712, GSM2645713, GSM2645714, GSM2645715). (e) Distribution of maintained and changed (with various types) TADs called from WT/KO cells when examined in 6KR/KO cells (left). Types of TAD changes are illustrated (right). TADs were identified with a window size of 250 kb. (f) Size distribution of chromatin interactions in WT/KO and 6KR/KO MCF7 cells. (g) Constitution of chromatin interactions for their anchorage at active promoters (indicated as prom), active enhancers (indicated as enh) or other regions in WT/KO and 6KR/KO MCF7 cells. (h) Aggregate peak analysis (APA) for MED1-occupied chromatin interactions identified from WT/KO and 6KR/KO cells with categorization according to their association with DEGs from 6KR/KO versus WT/KO cells (p=0.05 as cut-off). Z score (upper) and APA value (lower) are shown in each APA panel. Un-balanced matrices are used. (i) Distribution of chromatin interactions with both anchors occupied by MED1 for the anchors at high/mid/low ac groups of peaks (High Ac as top 25%, Low Ac as lowest 25%, Mid Ac as between). In all panels, chromatin interactions were identified in 10 kb bins as described in Methods.

**Supplementary Fig. 7.**
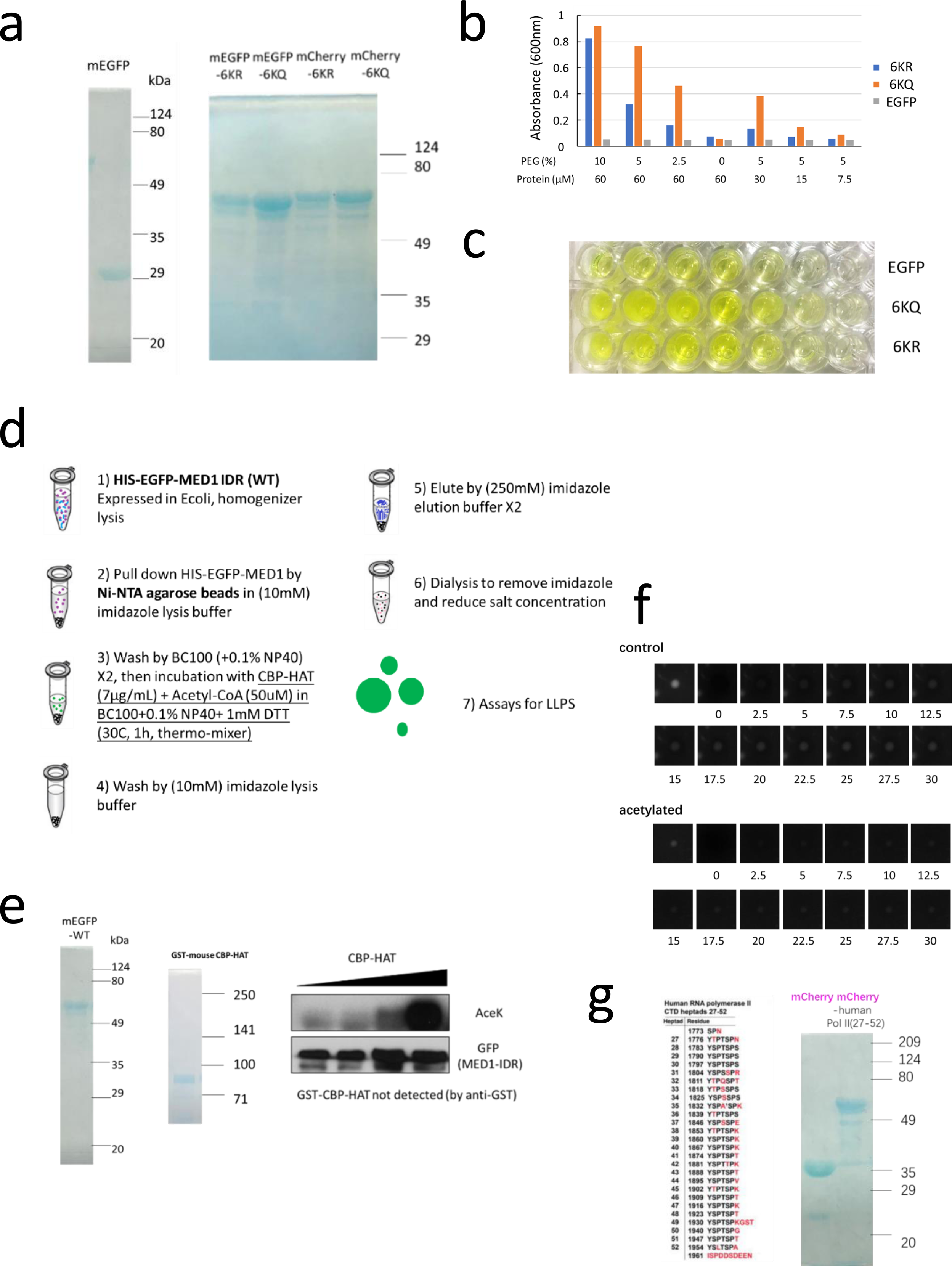
MED1 IDR acetylation mutation affects LLPS. (a) Colloidal blue staining for HIS-tagged mEGFP- and mCherry-fused MED1 IDRs (6KR or 6KQ, 963-1320 aa) that were expressed in *E.coli* and purified on Ni-NTA agarose beads. (b, c) Turbidity measured as absorbance of 600 nm demonstrating different LLPS propensities for 6KQ versus 6KR IDRs. Abs600 values (b) and plate images (c) are shown with PEG/protein concentrations indicated in the panel. (d) Workflow depicting that during purification, the bead-immobilized MED1 IDR was subjected to acetylation by incubation with CBP-HAT and acetyl-CoA, and then eluted and dialysed. (e) Colloidal blue staining for WT MED1 IDR fragment (fused to mEGFP) and CBP (GST-tagged) purified from *E.coli* (left), and immunoblotting confirming the enforced acetylation of MED1 IDR (right). (f) Faster recovery in FRAP of droplets formed by CBP-treated versus control WT MED1 IDR. Images for representative droplets are shown. (g) RNA Pol II CTD heptad sequences used in the Pol II co-condensation experiments (left) and colloidal blue staining of the mCherry-fused Pol II CTD heptads (right).

**Supplementary Fig. 8.**
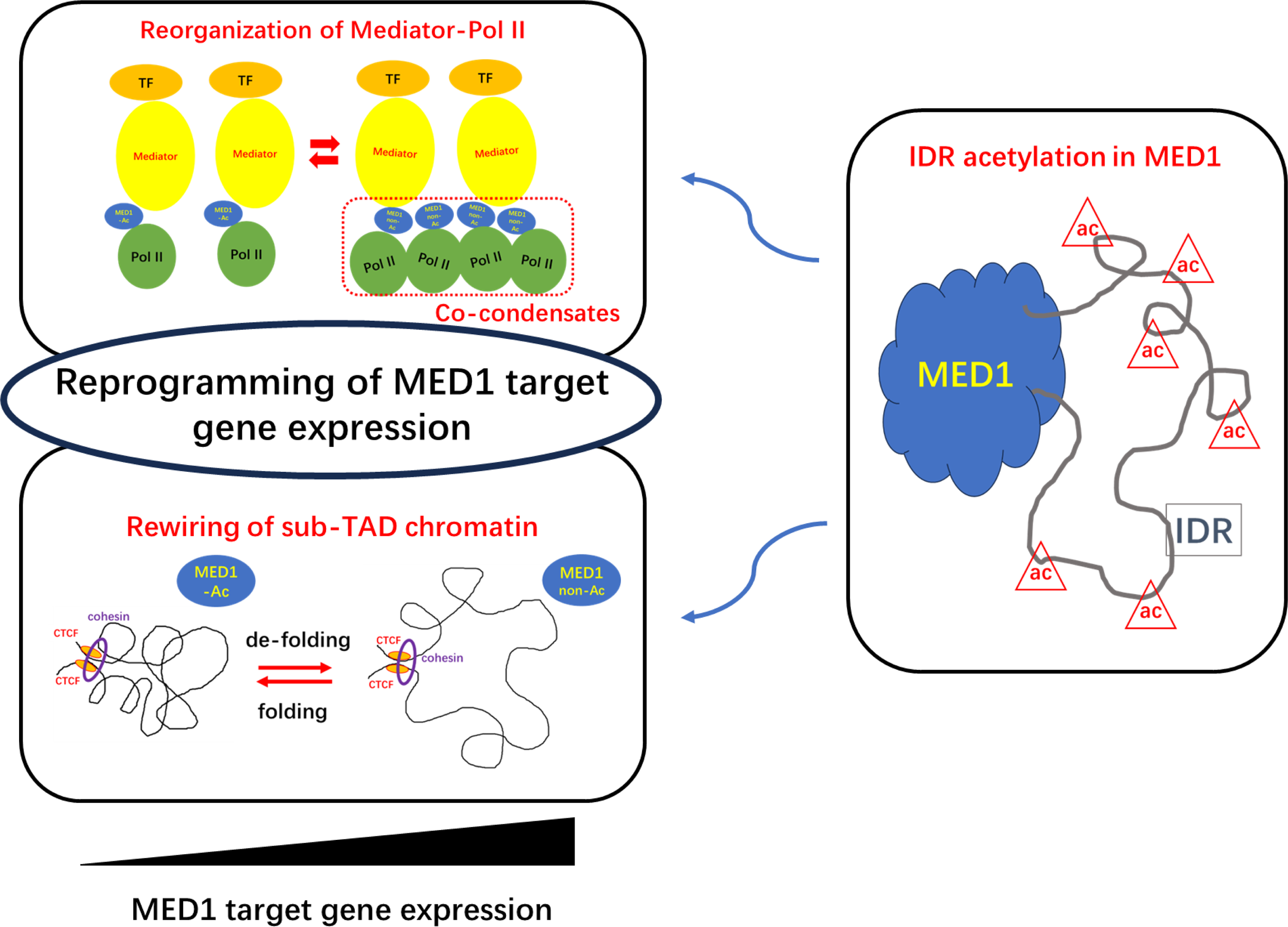
Graphic summary. Model of MED1 acetylation-controlled gene expression depicting that MED1 deacetylation results in (1) altered LLPS properties involved in MED1-Pol II interactions, which drives (either compositional or conformational) re-organization of transcription factor (TF)-MED1/Mediator-Pol II complexes, reflected as the assembly with higher MED1 and Pol II and lower MED17 (likely the entire Mediator core), while occupancy of transcription factors is unchanged; (2) unfolding of sub-TAD 3D chromatin architecture. Enhanced transcription activation of target genes is associated with such re-organized PICs and re-wired regional chromatin organizations.

## Methods

### 1. Cell culture

Human MCF7, T47D, MDA-MB231 cells (ATCC) were cultured in Dulbecco’s Modified Eagle Medium (Gibco) supplemented with 10% Fetal Bovine Serum (Gibco) in incubator at 37°C, with 5% CO2 and 95% humidity, and were tested regularly for Mycoplasma.

### 2. Stable transfection and pool selection, KO pool preparation

The two-plasmid lentivirus packaging system was employed to generate HEK293T or MCF7 cells with stable transgenic expression of FLAG-MED1 (WT or mutant), or HA-MED1 (WT or mutant), or FLAG-CAS9. The pLenti (CMV promoter) or pLX313 (EF1α promoter) constructs with inserted genes were co-transfected with vectors expressing lentiviral gag/pol (psPAX2) and vsvg (pMD2.G) genes in HEK293T cells to produce lentiviruses by TransIT-LT1 transfection reagent (Mirus). Supernatants containing virus particles were collected at 48 hours and 72 hours following transfection. After filtration through a 0.45 μm filter, collected virus-containing culture medium were used to infect HEK293T or MCF7 cells, facilitated by polybrene, followed by a selection in puromycin (Sigma-Aldrich) for 3 days or hygromycin (Sigma-Aldrich) for 7 days.

### 3. MED1 depletion in cells

To generate cell pool with MED1 gene depletion, cells with stable transgenic expression of FLAG-CAS9, and also HA-MED1 (with mutated gRNA targeting sites) in MED1 addback experiments, were infected by collected culture medium carrying lentivirus expressing gRNA targeting MED1 (or with mouse *Rosa26* sequence as non-targeting infection control) (sequence: see Table 1 attached below) where GFP was co-expressed. Cells were checked for near 100% GFP positive indicating the infection efficiency 3 days after infection and total protein from cells were extracted 4 or 5 days after infection for immunoblotting to confirm the depletion efficiency (by detecting MED1 expression in depletion-only Vs infection control cells) and expression level of ectopic MED1 (by detecting HA). Cell growth and colony formation assays were started with cells 5 days after infection, and RNA-seq, ChIP-seq, HiCAR samples were all collected at 5 days after infection.

**Table 1.**
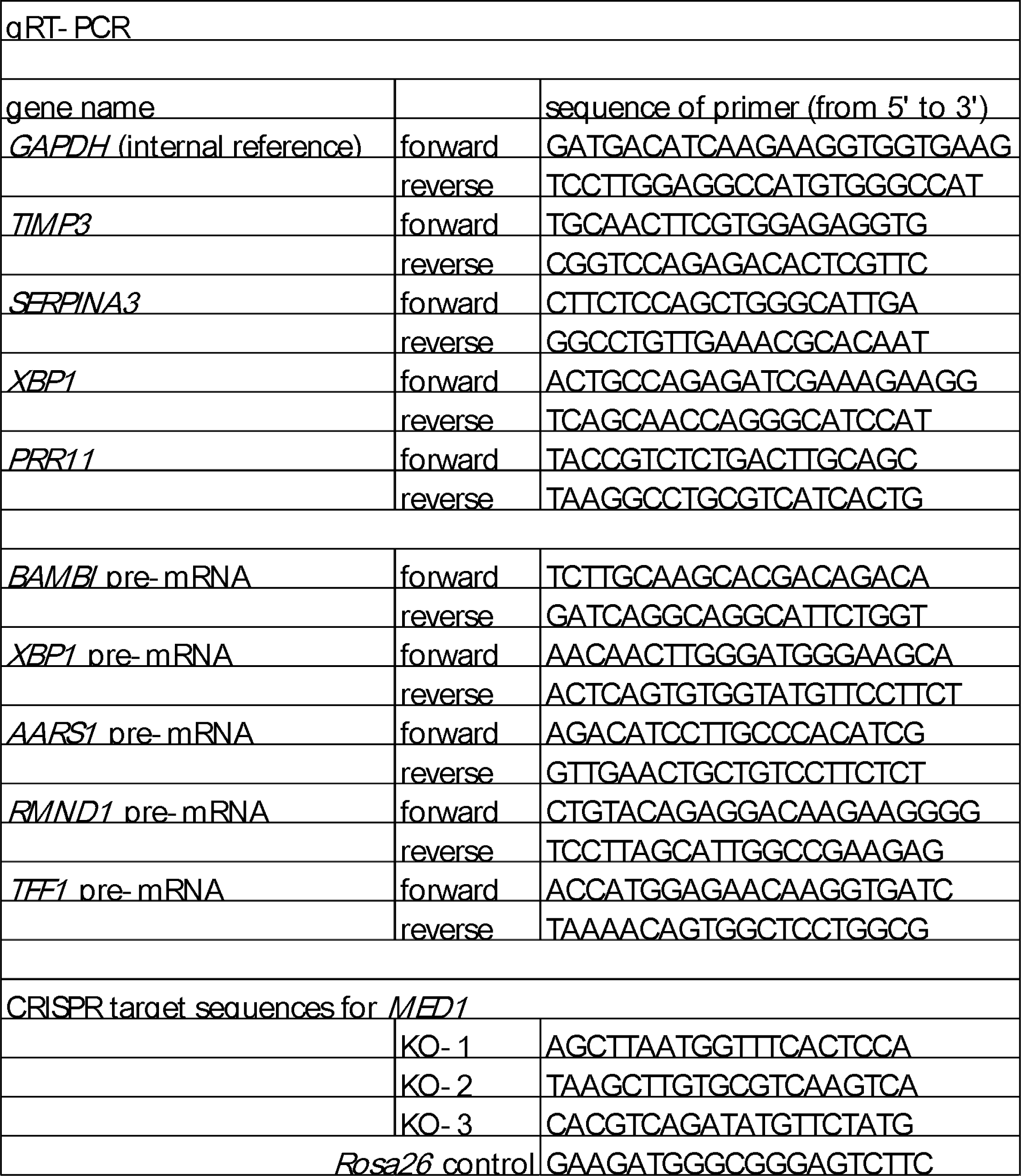
gRNA sequence, primer sequence for qPCR.

### 4. Cell growth – MTT and CellTiter Blue assays

To analyze cell growth, cells were digested and plated in 96-well plates at 6 repeat wells for each group in each experiment at a density of 500 or 1,000 cells per well in 100 μL culture medium. Cell growth was measured at day 0 and various time points after seeding. By using CellTiter Blue assay kit (Promega, G8081), 20 μL of CellTiter Blue reagent was added into each well. After 4 hours of incubation at 37°C, fluorescence was measured by SpectraMax M5 (Molecular Device) at 560/590 nm. Cell viability was calculated by subtracting the background fluorescence of culture medium and normalized against those at starting time (day 0). Cell growth was also monitored by traditional MTT assay. 20 μL Thiazolyl Blue Tetrazolium Bromide (Sigma-Aldrich) (5 mg/ml in PBS) was added into each well. After 4 hours of incubation at 37°C, culture medium was removed carefully and 200 μL DMSO was added into each dried well. After 45 min of mixing at room temperature, 560 nm absorbance was measured by SpectraMax M5.

### 5. Colony formation assay

To analyze colony formation, cells were digested and plated in 6-well plates at 3 repeat wells for each group at a density of 1,000 cells per well in 2 mL culture medium. After incubation for 1.5 to 2.5 weeks, culture medium was removed and colonies were stained with 0.1% crystal violet (Sigma-Aldrich) in 10% ethanol for 30 minutes.

### 6. Immunofluorescent staining

Cells were cultured in 6-well plates to the appropriate density and fixed with 4% paraformaldehyde (Sigma-Aldrich) for 15 min and then permeabilized with 0.02% Triton X-100 for 5 min at 4°C. After blocking with 0.5% BSA for 30 min, cells were incubated with mouse anti-FLAG antibodies (Sigma-Aldrich, F1804) diluted (1:2000) in 0.5% BSA solution for at 4°C overnight. After washing with TBST (0.05% Tween-20, 50 mM Tris-HCl, 150 mM NaCl, pH 7.6) for three times (5 min each), cells were incubated with Alexa Fluor 488 conjugated goat anti-mouse antibodies (Invitrogen, A11001) diluted (1:2000) in PBS for 1 hour. Then, cells were incubated with 4′,6-diamidino-2-phenylindole (Sigma-Aldrich) for 0.1 μg/ml for 5 min and then washed with PBS for three times (5 min each). Images were taken on a Nikon Eclipse Ti microscope and processed with NIS-elements BR software (Nikon).

### 7. Immunoprecipitation

Cells were lysed in RIPA lysis buffer (50 mM Tris, 300 mM NaCl, 1 mM EDTA, 1% Nonidet P-40, 0.1% SDS, 0.25% deoxycholate sodium, pH 8.0) with protease inhibitors (Roche). Insoluble fraction in lysate was spinned down at 12,000 rpm for 15 min at 4°C to discard. For Flag immunoprecipitations, the transferred supernatant was incubated with anti-Flag M2 affinity resin (Sigma-Aldrich, A2220) (20 uL per 1 mL lysate) at 4°C overnight. For immunoprecipitation of endogenous MED1, transferred supernatant was incubated with anti-MED1 antibody (Bethyl, A300-793A, 1:800) at 4°C overnight, and then added with Protein A sepharose CL-4B beads (Amersham, 17-0963-03) (20 uL per 1 mL lysate) for 4 hours more incubation. After three times of washing with ice-cold RIPA buffer, beads were re-suspended with Laemmli buffer (2% SDS, 50 mM Tris-HCl, 10% glycerol, 5% β-mercaptoethanol, 0.0005% Bromophenol Blue, pH 6.8) and boiled at 100°C for 10 minutes for immunoblotting.

### 8. IP-Mass spectrometry for acetylation site identification

To identify residues of MED1 with acetylation, the endogenous MED1 was immunoprecipitated from whole cell lysate of HEK293T by anti-MED1 antibody (Bethyl, A300-793A, 1:800) followed with protein A sepharose CL-4B beads (Amersham, 17-0963-03). The affinity capture sample was subjected to SDS-PAGE and stained by GelCode Blue stain reagent (Thermo Fisher scientific). The bands corresponding to MED1 were subjected to in-gel trypsin and chymotrypsin digestions. The LC-MS/MS analysis was performed on an Easy-nLC 1200 liquid chromatography system (Thermo Fisher Scientific) coupled to an Orbitrap Fusion Lumos mass spectrometer (Thermo Electron) equipped with a nano-electrospray ion source (Thermo Fisher Scientific). Tryptic peptides were dissolved in loading buffer (1% acetonitrile and 0.1% TFA), injected and separated using 15 cm x 100 μm ID column (C18, 1.9μm, 120Å) using a linear gradient created by mixing buffer B (80% acetonitrile in 0.1% formic acid) and Buffer A (0.1% formic acid). Eluted peptides were analyzed by data-dependent MS/MS acquisition. The scan range was set from m/z 375 to m/z 1500. Spectral data were searched against human protein UniProt database in Proteome Discoverer 1.4 suites with Mascot software (version 2.4, Matrix Science) and filtered for false discovery rate of <5%. The mass tolerance was set to be 7ppm for precursor, and it was set 0.0.02Da for the tolerance of product ions. Lysine acetylation (N-terminal), acetylation and phosphorylation were chosen as variable modifications.

### 9. Immunoblotting

Either whole cell lysate samples (post-confluent cells lysed directly in Laemmli buffer) or immunoprecipitated samples were resolved by SDS-PAGE (6% to 10%) and transferred onto methanol-treated immun-blot PVDF membrane (Bio-Rad). The membranes were incubated overnight at 4°C with corresponding primary antibodies (antibody information: see Table 2 attached). Membranes were then washed three times with TBST (0.05% Tween-20, 50 mM Tris-HCl, 150 mM NaCl, pH 7.6), incubated with ECL horseradish peroxidase-labeled anti-rabbit (Amersham NA934) or anti-mouse IgG (Amersham NA931) at dilution of 1:10,000 for 1 hour at room temperature, washed three times with TBST, exposed with mixed ECL Prime Peroxide Solution and ECL Prime Luminol Enhancer Solution (Amersham) and exposed to X-ray film for visualization.

**Table 2.**
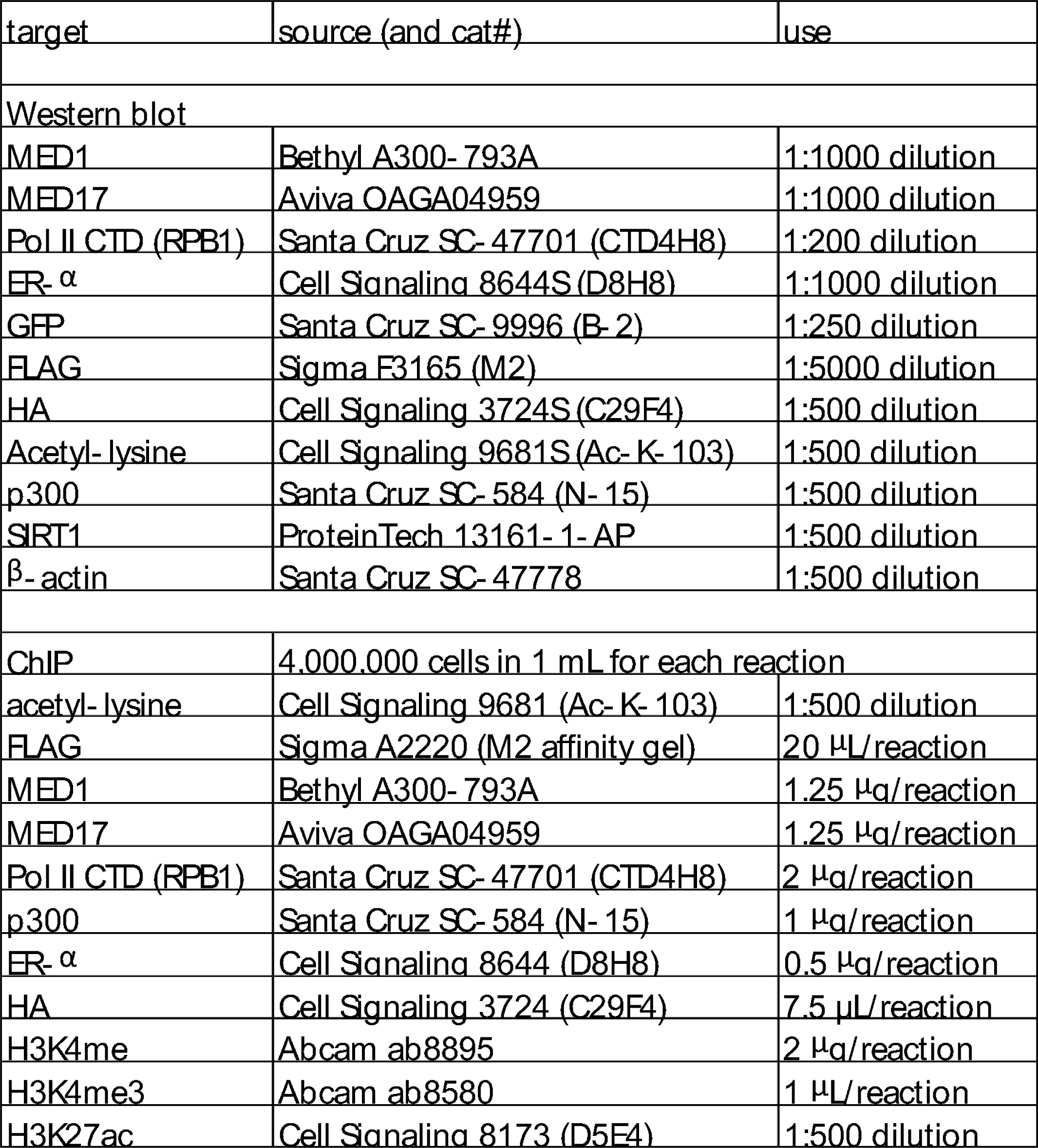
Antibody information.

### 10. Protein expression and purification from *E.coli*

The pET22b constructs of HIS-tagged mEGFP-MED1-IDR (WT or mutant) or mEGFP alone, the pET28a constructs of HIS-tagged mCherry-MED1-IDR (6KR/Q mutant), and the pET28c constructs of HIS-tagged mCherry-Pol II CTD (subcloned from addgene# 98678) or mCherry alone, and the pGEX2T constructs of GST-tagged CBP HAT domain (addgene #21093) were introduced into BL21 (DE3) *E.coli* cells. The transformants were grown at 37°C in LB medium (10 g/L tryptone, 5 g/L yeast extract, 10 g/L NaCl) untill the OD600 of medium reached 0.6. Protein expression in *E.coli* was stimulated by IPTG (Isopropyl β-D-1-thiogalactopyranoside, Sigma-Aldrich) at 1 mM at 16°C overnight. *E.coli* cells were spinned down and re-suspended in cold imidazole extraction buffer (50 mM Tris-HCl, 500 mM NaCl, 10 mM imidazole, pH 7.4) for HIS-tagged proteins, or BC100 buffer (20 mM Tris-HCl, 100 mM KCl, 0.2 mM EDTA, 20% glycerol, pH 7.9) for GST-tagged proteins with addition of 1% Triton X-100, 1 mM DTT and protease inhibitor cocktail (Roche), then lysed in a high-pressure homogenizer (EmulsiFlex-C5, Avestin), centrifuged at 12,000 rpm for 15 min at 4°C to remove insoluble debris. The fusion protein in the lysate was affinity-purified by Ni-NTA agarose beads (Qiagen, 30210) capturing HIS tag or by glutathione sepharose 4B beads (Amersham, 17-0756-01) capturing GST tag after 4 hours incubation at 4°C. After washing with cold imidazole extraction buffer or BC100 buffer for three times respectively, the HIS-tagged proteins were eluted by imidazole elution buffer (50 mM Tris-HCl, 500 mM NaCl, 250 mM imidazole, pH 7.4) for 10 min at 4°C for twice, and the GST-tagged proteins were eluted by glutathion elution buffer (50 mM Tris-HCl, 10 mM Reduced glutathione, 1 mM DTT, pH 8.0) for 10 min at room temperature for twice. Purity of recombinant proteins was verified by SDS-PAGE with staining by GelCode Blue stain reagent (Thermo Fisher scientific).

To prepare the acetylated WT MED1 IDR, the Ni-NTA agarose beads bound with HIS-tagged MED1 IDR (about 500 μL) were re-suspended with 500 μL BC100 buffer after washing with cold imidazole extraction buffer and GST-CBP was added to final concentration of 0, 0.708, 7.08, 70.8 μg/mL. After incubation at 30°C for 1 hour, the HIS-tagged MED1 IDR were eluted by imidazole elution buffer for 10 min at 4°C for twice. Purity of eluted proteins was verified by SDS-PAGE with staining by GelCode Blue stain reagent (Thermo Fisher scientific), and acetylation was verified by immunoblotting.

The eluted HIS-tagged MED1 IDR proteins and their controls (2-3 mL for each) were transferred into 15.5 mm size cellulose dialysis tube (BioDesign) and dialyzed with 1 L of dialysis buffer (50 mM Tris-HCl, 200 mM NaCl, 10% glycerol, 1 mM DTT, pH 7.4) for twice (overnight for the first time and 6 hours for the second time) to remove imidazole and reduce salt concentration. Dialyzed samples were concentrated in Ultracel centrifugal filters with 10K cellulose membrane (Amicon, UFC501096) at 8,000 rpm for 10-15 min. Concentration of purified proteins (with or without dialysis) was determined by Bradford method.

### 11. p300 purification

For recombinant p300, protocol was followed from a previous publication^55^. In brief, HIS-fused p300 was expressed from pFastBac1 vector in High-Five cells and purified on Ni-NTA agarose beads (Qiagen, 30210) in BC buffer (20 mM HEPES pH7.9, 1 mM EDTA, 10% glycerol and 1 mM DTT) with 100 mM NaCl and 300 mM imidazole. HIS-p300 was further purified on a Q sepharose column (HiTrap, GE Healthcare) in BC buffer and eluted between 100-600 mM NaCl. The purity of the protein preparation was assessed on gels, and the pure fractions (> 90%) were collected and flash frozen until use.

### 12. In vitro droplet formation and imaging

For droplet formation from a single IDR fragment with fluorescent tag, 10 μL dialysed and concentrated fragment in dialysis buffer (50 mM Tris-HCl, 200 mM NaCl, 10% glycerol, 1 mM DTT, pH 7.4) was mixed with 10 μL phase separation buffer (50 mM Tris-HCl, 10% glycerol, 1 mM DTT, pH 7.4) containing 20% polyethylene glycol (PEG) for 3 min at room temperature. For co-droplet tests, two types of dialysed and concentrated fragments were mixed (10 μL in total, with volume for each calculated based on concentration) together with 10 μL phase separation buffer containing 20% PEG for 3 min at room temperature. This resulted in the solution with 10% PEG and 100 mM NaCl. The different concentrations of PEG and NaCl, as well as the protein concentrations, were indicated in the text and figures.

After mixing, sample was loaded onto a glass slide and covered by a #1.5 coverslip. The slide was inverted, and droplets that settled on the coverslip were imaged using a DeltaVision Image Restoration Microscope (Leica) equipped with an inverted IX-71 microscope stand (Olympus), SSI light source (Insight), a CoolSNAP HQ monochrome CCD camera (Photometrics), a 60X/1.42 oil objective, and bandpass filter sets for detection of EGFP (525/50) and mCherry (632/60) fluorescence. Image acquisition was performed using Softworx software (Softworx), and images were saved as .dv files. Droplet size and fluorescent intensity measurements were performed with ImageJ. 3D reconstruction and surface area measurements for droplets were conducted using Imaris v9 software (Oxford Instruments) with its spot identification function. Huygens v22 software (SVI) was employed to measure the pixel-based correlation between mCherry and mEGFP signals in 3D reconstructed droplets using its co-localization wizard function and manual ROI selection. Deconvolved images were solely utilized for pixel-based correlation tests.

### 13. Fluorescence Recovery After Photobleaching (FRAP) for in vitro droplets

Following droplet formation and imaging setup as described above, FRAP assay was carried out on the same Deltavision microscope setup. One pre-bleach image was acquired using the conditions described above, followed by four pulses of a 488nm laser line for photobleaching target droplets. Postbleach images were collected for 30 seconds at 0.05 second intervals. Image acquisition was carried out using Softworx software. Fluorescent intensity measurements of slices of droplets were conducted with ImageJ for assessing the recovery of fluorescence.

### 14. In vitro turbidity assay

IDR fragment were mixed with equal volume of PEG-containing phase separation buffer in 96 well plates to prepare for samples in 50 mM Tris, 100 mM NaCl, 10% glycerol, 1 mM DTT (pH 7.4) but with varied PEG concentration and protein concentration. After mixing, the absorbance at 600nm was immediately measured by SpectraMax M5 (Molecular Device).

### 15. Purification of Mediator complex from HeLa-S

Nuclear extracts were prepared as described^56^ from a HeLa-derived cell line that stably expresses a FLAG-tagged human MED10. Mediator complex in the nuclear extract was affinity-purified by anti-FLAG M2 Affinity gel (Sigma-Aldrich, A2220) and washed extensively by BC100 buffer (20 mM Tris-HCl, 100 mM KCl, 0.2 mM EDTA, 20% glycerol, pH 7.9) with 0.1% NP-40, and then eluted by FLAG peptide (0.3 mg/mL) in BC100 for 3 times.

### 16. In vitro acetylation assay

Purified Mediator complex (0.2 μg) was incubated with or without p300 (0-0.2 μg, amount indicated in text/figures), and 50 μM Acetyl-CoA, in 12.5 μL of BC100 buffer (20 mM Tris-HCl, 100 mM KCl, 0.2 mM EDTA, 20% glycerol, pH 7.9) with addition of 0.1% NP-40 and 1 mM DTT. After 0-1 h (indicated in text/figures) at 30°C, samples were subjected to SDS-PAGE and immunoblotting.

### 17. RNA extraction and qRT-PCR, RNA-sequencing

Total RNA was extracted from post-confluent cells using TRIzol reagent (Ambion). The RNA template was converted into complementary cDNA using iScript Reverse Transcription SupemMix (Bio-Rad) with blend of oligo(dT) and random primers for complete RNA coverage. Real-time quantitative PCR reactions were performed using a 7300 Real-Time PCR System (Applied Biosystems) with QuantiTech SYBR Green PCR kit (Qiagen, 204141). The PCR protocol involved warming-up at 50°C for 2 min, pre-heating at 95°C for 10 min, denaturation at 95°C for 10 s and combined annealing and extension at 60°C for 1 min over 40 cycles. The melting curve was generated after these cycles (95°C for 10 s, 60°C for 1 min, 95°C for 15 s) to ensure that the amplification in each reaction was specific. GAPDH was taken as internal reference. Fold-changes of RNA levels were calculated using the ΔΔCt method. (Primer sequences used for qRT-PCR: see Table 1 attached)

For RNA-seq, total RNA was extracted using RNeasy Mini Kit (Qiagen, 74104) according to manufacturer’s instructions. Library of polyA-containing mRNA (350 bp size) was prepared using TruSeq Stranded mRNA Library Prep kit (Illumina) according to the manufacturer’s instructions. The library quality was evaluated using the Bioanalyzer and Illumina NovaSeq SP paired-end (100bp X 2) sequencing was performed by New York Genome Center.

### 18. Chromatin immunoprecipitation and ChIP-sequencing

Post-confluent cells were fixed with 1% formaldehyde for 10 min at room temparature, followed by quenching with 125 mM glycine and washed with cold TBS (20 mM Tris-HCl, 150 mM NaCl, pH 7.4) twice. Cells were lysed with the cell lysis buffer (10 mM Tris-HCl, 10 mM NaCl, 0.5% NP-40, pH7.5) by sitting on ice for 10 min. Cell lysates were spinned down and resuspended in 500 μL MNase digestion buffer (20 mM Tris-HCl, 15 mM NaCl, 60 mM KCl, 1 mM CaCl_2_, pH 7.5) with MNase (New England, M0247S, 0.25 U/1000 cells). After 20 min of incubation at 37°C with continuous mixing, digestion was stopped by addition of 500 μL of 2X STOP/ChIP buffer (100 mM Tris-HCl, 20 mM EDTA, 200 mM NaCl, 2% Triton X-100, 0.2% sodium deoxycholate, pH 8.1). Samples were sonicated for 7 cycles (30 s on/ 30 s off) with 40% amplitude (Branson Digital Sonifier) and then centrifuged at 12,000 rpm for 15 min at 4°C to remove insoluble debris. Apart from input samples (1% volume for each ChIP reaction) kept for reverse-crosslinking, lysates were incubated with antibodies (antibody information: see Table 2 attached) at 4°C overnight.

For sequential ChIP, the lysates were incubated with anti-FLAG M2 Affinity gel (Sigma-Aldrich, A2220) (20 μL per 1 mL lysate, equivalent to 4,000,000 cells) at 4°C overnight, then washed and re-suspended by STOP/ChIP buffer (50 mM Tris-HCl, 10 mM EDTA, 100 mM NaCl, 1% Triton X-100, 0.1% sodium deoxycholate, pH8.1) with addition of FLAG peptide (1 mg/mL) for 5 hours incubation at 4°C. After spinned down to discard the beads, eluate was further incubated with anti-acetyl-lysines (Cell Signaling, 9681) with 1:500 dilution at 4°C overnight.

Antibody-bound complexes were then captured by incubation with Protein A sepharose CL-4B beads (Amersham, 17-0963-03) for rabbit antibodies or Protein G sepharose 4B beads (Invitrogen, 101241) for mouse antibodies (both 20 μL per 1 mL lysate) for 4 h at 4 °C. The beads were washed sequentially with STOP/ChIP buffer (50 mM Tris-HCl, 10 mM EDTA, 100 mM NaCl, 1% Triton X-100, 0.1% sodium deoxycholate, pH 8.1), high salt buffer (50 mM Tris-HCl, 10 mM EDTA, 500 mM NaCl, 1% Triton X-100, 0.1% sodium deoxycholate, pH 8.1), LiCl_2_ buffer (10 mM Tris-HCl, 0.25 M LiCl_2_, 0.5% NP-40, 0.5% sodium deoxycholate, 1 mM EDTA, pH 8.0), and TE buffer (50 mM Tris-HCl, 10 mM EDTA, pH 8.0) (twice). Bound DNA was eluted and reverse-crosslinked at 65°C overnight in ChIP elution buffer (10 mM Tris-HCl, 150 mM NaCl, 10 mM EDTA, 5 mM DTT, 1% SDS, pH 8.0). After treatment with RNase A (0.2 mg/ml, 37°C, 1h) and proteinase K (2 mg/ml, 37°C, 2h), DNA used for sequencing was purified using a MiniElute PCR purification kit (Qiagen, 28004), and DNA used for qPCR was purified using a QIAquick PCR purification kit (Qiagen, 28106). DNA quality was assessed using an Agilent Bioanalyzer, and quantified using a Qubit Fluorometer (Invitrogen).

ChIP-seq libraries were generated using DNA SMART ChIP-seq kits (Takara Bio, 634865) according to the manufacturer’s instructions. After size selection (for 250-600 bp) by AMPure XP beads (Beckman Coulter, A63880), the library quality was evaluated using the Bioanalyzer and sequenced on Illumina NovaSeq SP (100 bp two ends) for sequential ChIP-seq and Illumina NextSeq High (75 bp single end) for all other ChIP samples by Genomics Resource Center in Rockefeller University.

### 19. HiCAR sequencing

The full protocol for the latest developed HiCAR workflow was published by Wei et al^57^. Collection of 400,000 cells were digested and fixed with 1% formaldehyde for 10 min, followed by quenching with 125 mM glycine and washed with cold TBS (20 mM Tris-HCl, 150 mM NaCl, pH 7.4) once. Cells were lysed by NPB buffer (PBS with 5% BSA, 0.2% NP-40, 1 mM DTT) and nuclei were spinned down and re-suspended with TB buffer (33 mM Tris-AC, 66 mM KAc, 10 mM MgAC2, 16% DMF, pH 7.8) where tagmentation by assembled Tn5 (Diagenode, C01070010) complex was performed. The nuclei were further lysed by 0.5% SDS at 62°C and 1.25% Triton X-100 at 37°C and then digested by CviQI (New England) followed by Tn5 adapter ligation which required help from splint oligo in RCB buffer (50 mM Tris-HCl, 50 mM NaCl, 0.2% SDS, pH 8.0). Next, cross-links were reversed and DNA was precipitated in glycogen-NaAC-Ethanol solution and re-suspended in Tris-HCl (10 mM, pH 8.0) where Nla III (New England) digestion was performed. AMPure XP beads (Beckman Coulter, A63880) was used for size selection and DNA at 1 ng/μL was circularized by T4 DNA ligase (New England) in Tris-HCl (10 mM, pH 8.0). Final library was amplified by Q5 polymerase (New England). The PCR condition: initial denaturation 98°C for 30 s, 12 cycles of denaturation 98°C for 10 s/ annealing 59°C for 30 s/ extension 72°C for 30 s, then final extension 72°C for 5 min. The size of library should range from 200 bp to 800 bp, so the DNA with smaller or larger size was removed by another round of AMPure XP beads purification. The sequencing was performed with Illumina NovaSeq SP (100 bp two ends) by Genomics Resource Center in Rockefeller University. All PCR primer/oligo sequences were reported by Wei, et al^57^.

### 20. Bioinformatic data analysis

#### 20.1. RNA-seq analysis

Transcript abundance was determined from FASTQ files using Salmon (v0.8.1) and the GENCODE reference transcript sequences^58^. Transcript counts were imported into R with the tximport R Bioconductor package (v1.8.0), and differential gene expression was performed with the DESeq2 R Bioconductor package (v1.20.0) ^59^. Normalized counts were retrieved from the DESeq2 results and z-scores for the indicated gene sets were visualized with heatmaps generated using the pheatmap R package (v 2.8.0) (R. Kolde. Pheatmap: Pretty Heatmaps. R package version 1.0.12). For GSEA, the Wald statistic from the DESeq2 analysis was used to generate the ranked list of genes. The clusterProfiler R Bioconductor package (v4.0.5) was then used to perform GSEA to determine the enrichment of the indicated gene lists within this ranked list ^60,61^. The enrichKEGG function from clusterProfiler was used for over representation analysis (Fisher test) of KEGG gene lists in differentially expressed gene sets. The enrichplot R Bioconductor package (v1.12.3) was used to visualize the GSEA result (G Yu. Enrichplot: Visualization of Functional Enrichment Result. R package version 1.12.3). Volcano plots showing log2 fold-change versus log10 of the adjusted p-value and the plot showing the correlation between RNA seq data sets were generated using the ggplot2 R package (v3.3.5) (H. Wickham. Ggplot2: Elegant Graphics for Data Analysis. 2016). When genes were divided into categories based on the degree of their gene expression changes, those groups are defined as follows: (1) strong up: fold-change is greater than 2 and adjusted p-value is less than 0.05; (2) mild up: fold-change is between 0 to 2 and adjusted p-value is less than 0.05; (3) strong down: fold-change is less than 2 and adjusted p-value is less than 0.05; (4) mild down: fold-change is between 0 to 2 and adjusted p-value is less than 0.05; (5) no change: adjusted p-value is greater than 0.05.

#### 20.2. ChIP-seq analysis

ChIP-seq reads were aligned using the Rsubread R Bioconductor package (v1.30.6) and predicted fragment lengths were calculated by the ChIPQC R Bioconductor package (v1.16.2)^62,63^. Normalized (reads at each loci per million mapped reads), fragment-extended signal bigWigs were created using the rtracklayer R Bioconductor package (v1.40.6), and peaks were called using MACS2 (v2.1.1)^64,65^. Range-based heatmaps, line plots, and boxplots showing normalized signal over genomic regions were generated using the profileplyr R Bioconductor package (v1.8.1) and ggplot2 (Barrows D, Carroll T. profileplyr: Visualization and annotation of read signal over genomic ranges with profileplyr. R Package Version 1.2.0). Further normalization of bigwig files (normalized either to input or knockout cells) was performed using the deeptools (v3.5.1) bigwigCompare function with the ‘operation’ argument set to log2^66^. Correlation of signal from ChIP-seq replicates was performed using the deeptools ‘multiBigwigSummary’ function with the‘binSize’ argument set to 5000, followed by the ‘plotCorrelation’ function with using a Pearson correlation. Annotation of peaks to genes was done with the rGREAT R Bioconductor package (v1.24.0). Any regions included in the ENCODE blacklisted regions of the genome were excluded from all region-specific analyses^67^.

#### 20.3. HiCAR analyses

Fastq files were run through the Nextflow HiCAR pipeline, using default settings (https://nf-co.re/hicar) (nf-core/hicar Release 1.0.0. Zenodo. https://doi.org/10.5281/zenodo.6515313). Cooler files from the Nextflow pipeline were visualized as heatmaps at 2.5 Mb, 50 kb, and 5 kb resolution using the HiGlass tool ^68^. To determine quantitative changes of read counts in interactions between WT/KO and 6KR/KO MED1 cells, data from cooler files were extracted at 10K resolution using the ‘cooler dump’ function from the cooler command line interface (v0.9.1) ^69^. The HiCcompare R Bioconductor package was used to read unbalanced count data into R, adjust for sequencing depth, apply loess normalization, and determine fold-changes (v1.12.0)^70^. To perform overlaps and annotations of loops called with the pipeline, contact data was converted to a GInteractions object from the InteractionSet R Bioconductor package for downstream analysis (v1.20.0)^71^. Overlaps with active enhancers, active promoters, or acetylated MED1 peak sets were assessed using the GRanges objects for each anchor. The Venn diagram showing overlap between the WT/KO and 6KR/KO samples was generated with overlaps determined using the findOverlaps function from the InteractionSet R Bioconductor package.

Aggregate peak analysis was performed on unbalanced contact data with the GENOVA R package (v1.0.1)^72^. Prior to APA analysis, anchors for each loop were annotated to genes using the rGREAT R Bioconductor package and it was determined whether either anchor of a loop was annotated by a gene that significantly changed in the RNA-seq WT/KO vs 6KR/KO MED1 dataset. Loops annotated by both a gene that went up and a gene that went down were not included in the APA analysis. Zoomed-in heatmaps around the TFF1 gene were generated with the GENOVA R package.

TADs were determined using the balanced 10kb resolution cool file with the ‘insulation’ function from the cooltools command line interface (v0.5.4) (open2c/cooltools: v0.5.4 (v0.5.4). Zenodo. https://doi.org/10.5281/zenodo.7563392). For TAD calling, the Li thresholding method was used with a window size of 250kb. Aggregate TAD analysis was performed on with the unbalanced cool file at 10kb resolution using the GENOVA R package.

To compare the HiCAR data to published HiC from MCF7, MDA-MB-231, or THP-1 cells (GEO accession #’s: GSE99541, GSE223785, GSE226673, respectively), hicrep (python version 0.2.6) was used to generate SCC scores from cooler files with a bin size of 10kb^38^.

